# Dynamic reconfiguration of functional brain networks during working memory training

**DOI:** 10.1101/685487

**Authors:** Karolina Finc, Kamil Bonna, Xiaosong He, David M. Lydon-Staley, Simone Kühn, Włodzisław Duch, Danielle S. Bassett

## Abstract

The functional network of the brain continually adapts to changing environmental demands. The consequence of behavioral automation for task-related functional network architecture remains far from understood. We investigated the neural reflections of behavioral automation as participants mastered a dual n-back task. In four fMRI scans equally spanning a 6-week training period, we assessed brain network modularity, a substrate for adaptation in biological systems. We found that whole-brain modularity steadily increased during training for both conditions of the dual n-back task. In a dynamic analysis, we found that the autonomy of the default mode system and integration among task-positive systems were modulated by training. The automation of the n-back task through training resulted in non-linear changes in integration between the fronto-parietal and default mode systems, and integration with the subcortical system. Our findings suggest that the automation of a cognitively demanding task may result in more segregated network organization.

## INTRODUCTION

The brain constantly adjusts its architecture to meet the demands of the ever-changing environment. Such neural adaptation spans multiple time scales, being observed over seconds to minutes during task performance^1–5^, over days to weeks during learning^6–8^, and over years during development^9^. Like many other complex biological systems, the adaptability of the brain is supported by its modular structure^10^. Intuitively, modularity allows for dynamic switching between states of segregated and integrated information processing, whose balance is constantly adjusted to meet the requirements of our cognitive faculties^11,12^. Understanding the patterns of these adjustments and determining the rules that explicate their relation to human behavior is one of the most important challenges for cognitive neuroscience.

It is hypothesized that simple, highly automated sensorimotor tasks can be maintained by a highly segregated brain organization, while more complex and cognitively demanding tasks require integration between multiple subnetworks^13^. Indeed, switching from a segregated to a more costly integrated network architecture is consistently reported as human participants transition to challenging tasks with heavy cognitive load^1–5^; in contrast, network organization during simple motor tasks remains highly segregated^3,4^. Whether shifts towards network integration depend on the level of task complexity or on the level of task automation remains to be delineated^12^. Is it possible that a complex, but fully automated task, can be performed without the need for costly network integration?

Longitudinal studies, during which participants are scanned multiple times while mastering a specific task, can shed light on patterns of network adaptation related to learning and task automation^12^. For example, Bassett et al.^7^ showed that training on a visuomotor task over the course of 6 weeks leads to increased autonomy between task-relevant subnetworks in motor and visual cortices. In another study, Mohr et al.^8^ found increased segregation of the default mode system after short-term visuomotor training. Collectively, these findings suggest that an increase in network segregation and a decrease in integration may constitute a natural consequence of task automation. However, these results refer to the training of simple motor tasks, which do not require extensive network integration, in contrast to complex tasks involving higher order cognitive functions such as cognitive control^12^. The consequence of complex cognitive task automation on the balance between network segregation and network integration remains unknown.

In the present study, we investigated whether mastering a demanding working memory task affects the balance between network segregation and integration during task performance. Does effortless performance of the demanding cognitive task lead to the same increase in network segregation that is characteristic of simple motor tasks^3,8^? Is the breakdown of network segregation during the changing demands of the cognitive task still necessary when the cognitive task is automated? Finally, do we observe stronger separation of subnetworks relevant to cognitive control when tracking dynamical brain network reorganization throughout the course of training? To address these questions, participants underwent four functional magnetic resonance imaging (fMRI) scans while performing an adaptive dual n-back task taxing working memory over a 6-week training period. The dual n-back task consisted of visuospatial and auditory tasks that were performed simultaneously^14^. In the visuospatial portion of the task, participants had to determine whether the location of the stimulus square presented on the screen was the same as the location of the square n-back times in the sequence; in the auditory portion of the task participants had to determine whether the heard consonant was the same as the consonant they heard n-back times in the sequence. To ensure that participants mastered the task due to training, and not simply due to a repeated exposure to the task, we compared their performance to an active control group. While participants from both the experimental and the control groups performed the same version of the dual n-back task, with interleaved 1-back and 2-back blocks, inside the fMRI scanner, only the experimental group trained their working memory using an adaptive version of the task in 18 training sessions outside the scanner. We examined network reconfiguration using static functional network measures to distinguish distinct task conditions, and using dynamic network measures to study fluctuations of network topology across short task blocks.

First, we investigated global changes in network segregation (modularity) across different task conditions as compared to rest. In line with the aforementioned research, we expected modularity to decrease during dual n-back task performance compared to rest, and also to decrease as the demands of the n-back task increased. We also hypothesized that over the course of training network segregation during the n-back task would increase, and the extent of demand-related modularity change would decrease. In the systems relevant to working memory performance – the fronto-parietal and the default mode systems^15^ – we expected an increase in autonomy throughout the course of training. To verify this hypothesis, we utilized previously developed dynamic network methods^7^ to assess the recruitment and integration of the default mode and fronto-parietal systems. Finally, we expected that changes in network architecture would correspond to the level of task automation and training progress.

Our results demonstrate that adult human brain functional networks not only reorganize during a working memory task, but also can be modulated by the level of expertise in the task. After working memory training, brain networks are more segregated. The increase in segregation is visible at the whole-brain level for static networks, and also evidenced by an increased segregation of the default mode and task-positive systems when considering dynamic changes in network organization. Automation of the working memory task is accompanied by non-linear changes in coupling between the default mode and fronto-parietal systems and engagement of the subcortical system. Together, these results shed new light on the mechanisms underlying brain network reorganization accompanying the automation of performance on cognitively demanding tasks.

## RESULTS

### Behavioral changes during training

Behavioral improvement in the task can either occur as a result of training or occur in response to repeated exposure to a task across multiple scanning sessions. To distinguish the effect of intensive working memory practice and task automation from the effect of repeated exposure, we employed an active control group. When participants from the experimental group underwent the challenging, adaptive, dual n-back working memory training, participants from the control group performed a single, non-adaptive, 1-back working memory task (Figure 1).

**Figure 1:**
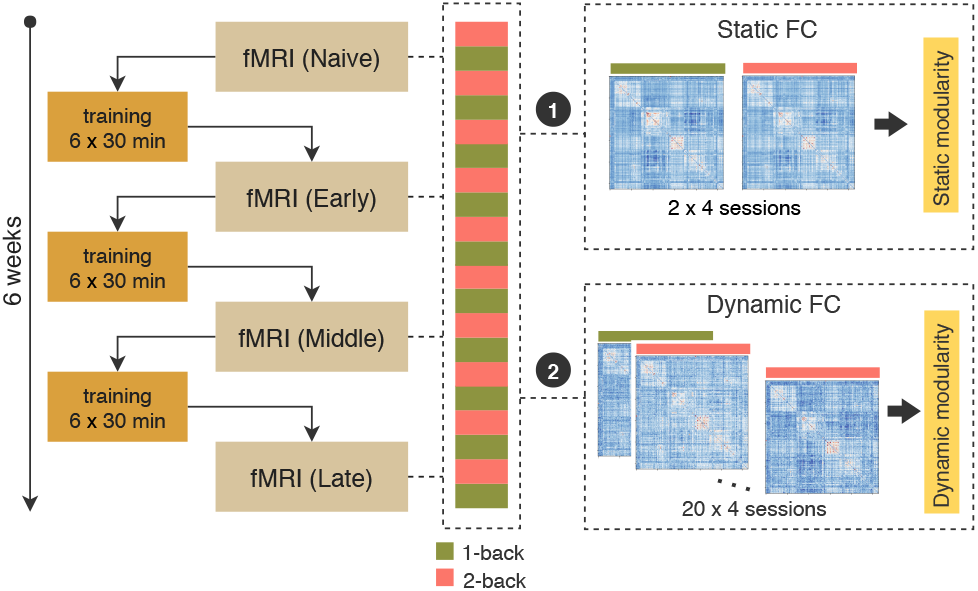
Study design. *(Left)* The dual n-back working memory task was performed in the scanner on the first day of the experiment (Naive), after 2 weeks of training (Early), after 4 weeks of training (Middle), and after 6 weeks of training (Late). *(Right)* We investigated (**1**) changes in static modularity across task conditions (1-back versus 2-back) and (**2**) dynamic fluctuations in network community structure from block to block.

The dual n-back task (1-back and 2-back conditions) was performed in the scanner on the first day of the experiment (Naive), after two weeks of training (Early), after four weeks of training (Middle), and after six weeks of training (Late). We measured participant performance as a *d*′, a measure based on signal detection theory that takes into account both response sensitivity and response bias^16^ (see Methods). Better cognitive performance is characterized by higher values of *d*′. We expected that participants from the experimental group would exhibit a substantial increase of *d*′ during training, particularly for the 2-back condition in comparison to the 1-back condition, the latter being easy to master even without extensive training.

Using multilevel modelling (see Methods), we found that participants had significantly different *d*′ depending on the training stage (Naive, Early, Middle, Late), condition (1-back vs. 2-back), and group (Experimental vs. Control). Specifically, we found a significant session × condition × group interaction (*χ*^2^(3) = 9.39, *p* = 0.02; Figure 2). The greatest improvement was observed in the experimental group when comparing ‘Naive’ to ‘Late’ training phases during the 2-back condition (mean 43.2% *d*′ improvement; Bonferroni-corrected, *t*(20) = −9.17, *p* < 0.0001). For comparison, the control group exhibited a 24.3% increase in *d*′ during the 2-back condition (Bonferroni-corrected, *t*(20) = −6.45, *p* < 0.0001). The increase in *d*′ was significantly larger for the experimental group than for the control group (Bonferroni-corrected, *t*(20) = −4.12, *p* = 0.0004; Figure 2d). In the 1-back condition, the experimental group displayed a 12.2% increase in *d*′ (Bonferroni-corrected, *t*(20) = −3.18, *p* = 0.02); no improvement was found in the control group (Bonferroni-corrected, *t*(22) = −1.91, *p* = 0.28) (see Figure 2c). The change in *d*′ during the 1-back condition did not differ between the two groups (*t*(39.64) = −0.52, *p* = 0.47). Interestingly, in the experimental group we observed no significant difference in performance between the 1-back condition and the 2-back condition after training (*t*(20) = 0.02, *p* = 0.98), while in the control group, the difference in performance between conditions remained substantial (Bonferroni-corrected, *t*(20) = 4.91, *p* < 0.0016). This finding suggests that the 2-back condition, which was much more effortful before training (‘Naive’ phase), was performed effortlessly after training, at the same level as the 1-back task.

**Figure 2:**
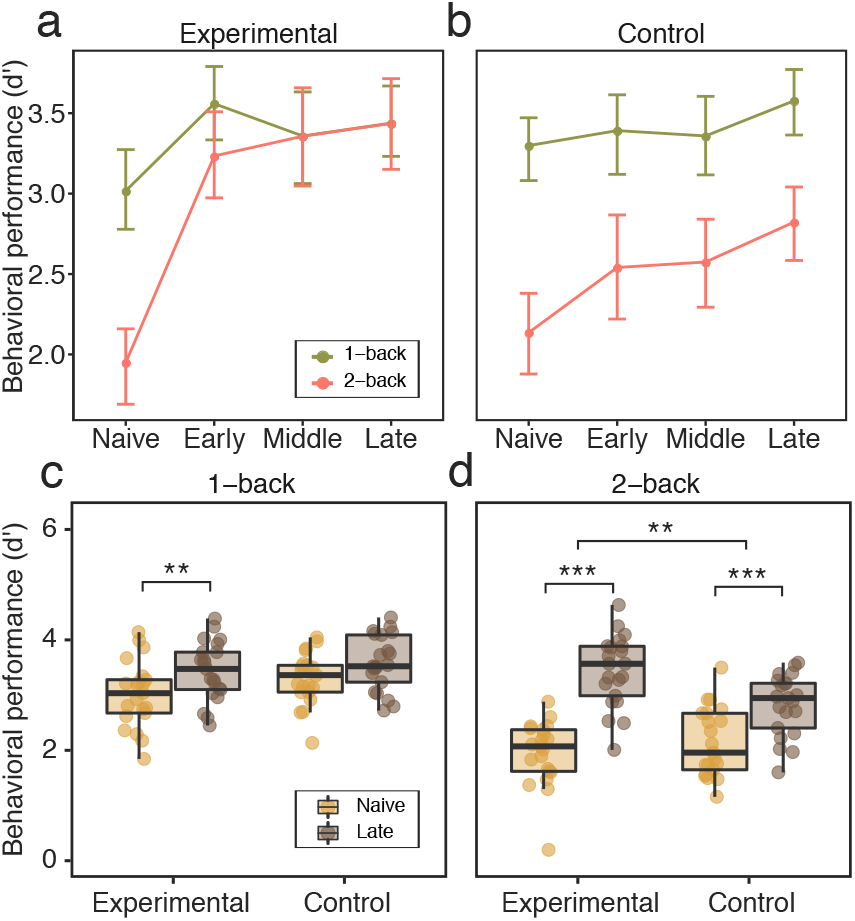
Behavioral performance modulated by training. (**a, b**) Line plots representing mean behavioral performance measured as *d*′, calculated for all training phases (Naive, Early, Middle, Late), dual n-back conditions (1-back and 2-back), and groups ((**a**) experimental and (**b**) control). We found a significant interaction effect between session, condition, and group. After training, the experimental group exhibited no difference in behavioral performance between the 1-back and 2-back conditions. (**c**) No significant difference between groups was found for *d*′ reduction (from Naive to Late sessions) during the 1-back task condition. (**d**) The experimental group showed a significant reduction in *d*′ compared to the control group during the challenging 2-back condition. Error bars represent 95% confidence intervals. *** *p* < 0.001 Bonferroni corrected; ** *p* < 0.05 Bonferroni corrected. Source data are provided as a Source Data file.

In sum, the results demonstrate that the experimental group gradually improved in behavioral performance measured during the fMRI scanning sessions, and that this improvement was significantly greater than the corresponding effect in the control group. We also replicated these findings using an alternative measure of behavior, penalized reaction time (pRT) which incorporates a measure of accuracy (see Supplementary Figure 3 and Supplementary Methods).

### Whole-brain network modularity changes

To establish whether complex working memory task training leads to increased network segregation at the whole-brain level, we investigated network modularity during different sessions and load conditions. Here, we employed a common community detection algorithm known as modularity maximization^17^, which we implemented using a Louvain-like locally greedy algorithm. The modularity quality function to be optimized encodes the extent to which the network can be divided into non-overlapping *communities*. Intuitively, a community is a group of densely interconnected nodes with sparse connections to the rest of the network^17^. Modularity is a relatively simple measure of segregation, with high values indicating greater segregation of the brain into non-overlapping communities and low values indicating lesser segregation. Because modularity depends upon the network’s total connectivity strength, we normalized each modularity score by dividing it by the mean of the corresponding null distribution calculated on a set of randomly rewired versions of the original networks^18^ (see Methods for details).

Functional network modularity may vary depending on the difficulty of the task. Several studies have reported a reduction in modular structure during demanding n-back conditions^2,3,5^. Here, we first investigated the differences between the high-demand 2-back condition and the low-demand 1-back condition as compared to a baseline resting state scan acquired during the first session (‘Naive’) for all subjects. Using multilevel modeling we found a significant main effect of condition (*χ*^2^(2) = 84.13, *p* < 0.00001). Planned contrast analysis revealed that network modularity during the dual n-back task was lower than network modularity during the resting state (*β* = −0.20, *t*(88) = −11.37, *p* < 0.00001). Furthermore, modularity was significantly reduced during the 2-back condition relative to the 1-back condition (*β* = −0.08, *t*(296) = −2.60, *p* = 0.01; Figure 3). We note that the results reported here use a functional brain parcellation composed of 264 regions of interests provided by Power et al.^19^; in robustness tests, we performed the same analyses using the Schaefer parcellation, and obtained similar results (see Supplementary Figure 16).

**Figure 3:**
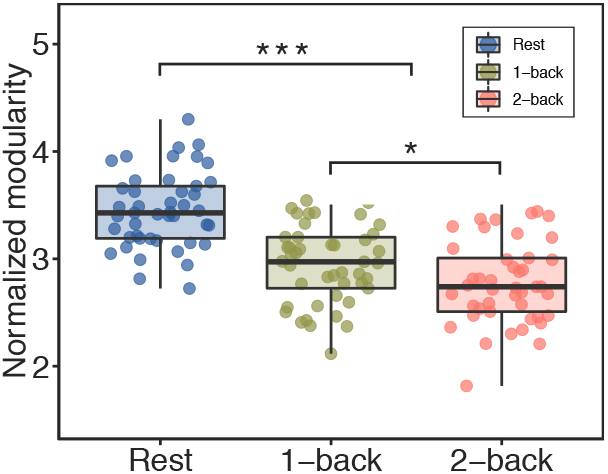
Modularity differences between resting and dual n-back task conditions in ‘Naive’ session. Whole-brain modularity was higher during the resting state than during the dual n-back task, and decreased as demands heightened from the 1-back to the 2-back condition. Error bars represent 95% confidence intervals. \*\*\**p* < 0.001, \**p* < 0.05. Source data are provided as a Source Data file.

The modularity of functional brain network architecture decreases appreciably during challenging task conditions, but is the breakdown in modularity still present when the demanding task is mastered? To address this question, we tested whether modularity during the dual n-back task changed depending on the session, task condition, and group. Using a multilevel model (see Methods), we found a significant main effect of session (*χ*^2^(2) = 19.40, *p* = 0.0002) and of group (*χ*^2^(1) = 6.62, *p* = 0.01). However, the experimental and control groups did not differ by session (*χ*^2^(1) = 1.44, *p* = 0.69), nor did we observe a significant session by condition interaction (*χ*^2^(1) = 1.50, *p* = 0.68). A planned contrast comparison showed that participants’ whole-brain functional network modularity significantly increased from ‘Naive’ to ‘Middle’ sessions (*β*= 0.15, *t*(114) = 2.61, *p* = 0.01) and from ‘Naive’ to ‘Late’ sessions (*β*= 0.24, *t*(114) = 4.05, *p* = 0.0001; Figure 4ab). The experimental group showed a higher network modularity (M = 3.09) than the control group (M = 2.87). To summarize, we showed that the modularity of the functional brain network generally increased during the training period. However, the degree to which modularity changed between load conditions remained stable. Groups did not differ significantly in the change of modularity. These results suggest that the functional brain network shifts towards a more segregated organization as a result of behavioral improvement after training and also after repeated exposure to the task. Although network modularity increased to a similar extent in both conditions, the demand-dependent change in modularity remained stable. One could interpret these results as suggesting that a general increase in modularity reflects the fact that less expensive information processing is required within segregated brain subsystems after training of the complex task.

**Figure 4:**
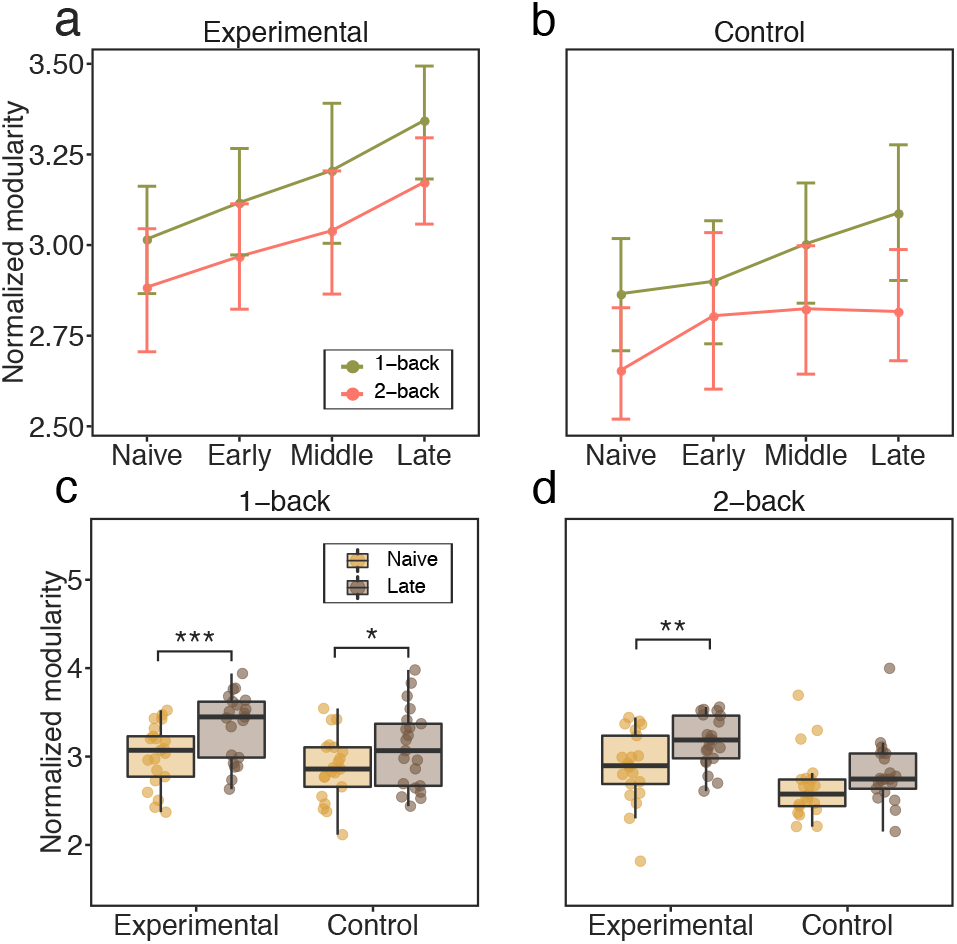
Modularity differences across task, sessions, and groups. (**a, b**) Line plots representing the mean values of modularity for each scanning session (Naive, Late, Middle, Late) and condition, separately for (**a**) the experimental group and (**b**) the control group. (**c, d**) Modularity changes from ‘Naive’ to ‘Late’ sessions for the 1-back condition and the 2-back condition. Error bars represent 95% confidence intervals. *** *p* < 0.01 Bonferroni corrected; ** *p* < 0.05 Bonferroni corrected, * *p* < 0.05 uncorrected. Source data are provided as a Source Data file.

To further explore the changes in modularity that might be specific to each group and condition, we performed additional analyses comparing modularity measured before and after training (Figure 4cd). Specifically, we employed separate paired *t*-tests to investigate differences in modularity for each group and condition between ‘Naive’ and ‘Late’ sessions. We found a significant increase of modularity in the experimental group in the 1-back condition (Bonferroni-corrected, *t*(20) = −3.66, *p* = 0.006) and in the 2-back condition (Bonferroni-corrected, *t*(20) = −3.33, *p* = 0.013). The increase in modularity observed in the control group was not significant for either the 1-back condition (Bonferroni-corrected, *t*(20) = −2.35, *p* = 0.11) or the 2-back condition (Bonferroni-corrected, *t*(20) = −1.88, *p* = 0.28). The change of modularity from ‘Naive’ to ‘Late’ sessions did not significantly differ between groups for the 1-back condition (*t*(39.88) = −0.80, *p* = 0.42) or for the 2-back condition (*t*(39.99) = −1.05, *p* = 0.30). These results indicate that the experimental group displays increased network modularity for both task conditions when moving from ‘Naive’ to ‘Late’ sessions, suggesting that network segregation may be a consequence of the 6-week working memory training. While the same effect was not present in the control group, we did not observe a significant group × session interaction, and therefore further work is needed to inform our conclusions.

Behavioral gains resulting from working memory training differed across participants, suggesting the existence of individual differences in learning capabilities. Therefore, we also tested whether the increase of modularity observed during the 2-back condition in the experimental group was correlated with behavioral performance after training as measured by a decrease in *d*′. However, we did not find a significant relationship between these two variables (Pearson’s correlation coefficient *r* = 0.08, *p* = 0.71; Supplementary Figure 20). This finding suggests that the change of modularity is a general consequence of training and may not reflect individual differences in behavioral improvement.

Our results confirmed the existence of a decrease in modularity during increased cognitive demands. However, changes in modularity during training were not different across conditions or experimental groups. A significant increase in modularity from ‘Naive’ to ‘Late’ sessions was found for the 1-back and 2-back conditions for the experimental group, which suggests the enhancement of network segregation associated with task automation.

### Dynamic reorganization of large-scale systems

The modular architecture of functional brain networks is not static but instead can fluctuate appreciably over task blocks. Here, we used a dynamic network approach to answer the question of whether large-scale brain systems change in their fluctuating patterns of expression during training. Based on a previous study of motor sequence learning^7^, we expected that systems relevant to working memory – the fronto-parietal and the default mode – would become more autonomous over the 6 weeks of working memory training (Figure 5a). To formally test our expectation, we investigated the dynamic reconfiguration of the network’s modular structure as subjects switched between blocks of the dual n-back task. Pooling across conditions and sessions, we constructed a multilayer network model of the data in which each block corresponds to a unique layer, each region corresponds to a node, and each functional connection corresponds to an edge. We then employed a multilayer community detection algorithm that estimates each node’s module assignment in each network layer^20^. The presence of fluctuations in community structure across task blocks is indicated by variable assignments of nodes to modules across layers. For each subject and session, we summarized these data in a module allegiance matrix **P**, where each element *P_ij_* represents a proportion of blocks for which node *i* and node *j* were assigned to the same module. We also applied a normalization to allegiance matrices, to remove any potential bias introduced by differences in the number of nodes within each subsystem. Following the functional cartography framework described by Mattar et al.^21^, we used **P** to calculate the recruitment of all 13 large-scale systems, as well as the pairwise integration among them (see Methods for details). We selected these measures to maintain consistency with the methodology used in a previous study on the effects of motor sequence training on the dynamics of functional brain networks^7^. Recruitment is defined for each system separately, while integration is calculated for pairs of systems. Intuitively, high recruitment indicates that nodes of the system are consistently assigned to the same module across different layers; this consistency reflects the non-random nature of brain dynamics in which a functional module is persistently recruited for a task. High integration indicates that pairs of nodes (where one region of the pair is located in one system and the other region of the pair is located in the other system) are frequently classified in the same module across layers). We used a multilevel model to test whether recruitment and integration coefficients differed between scanning sessions and experimental groups.

**Figure 5:**
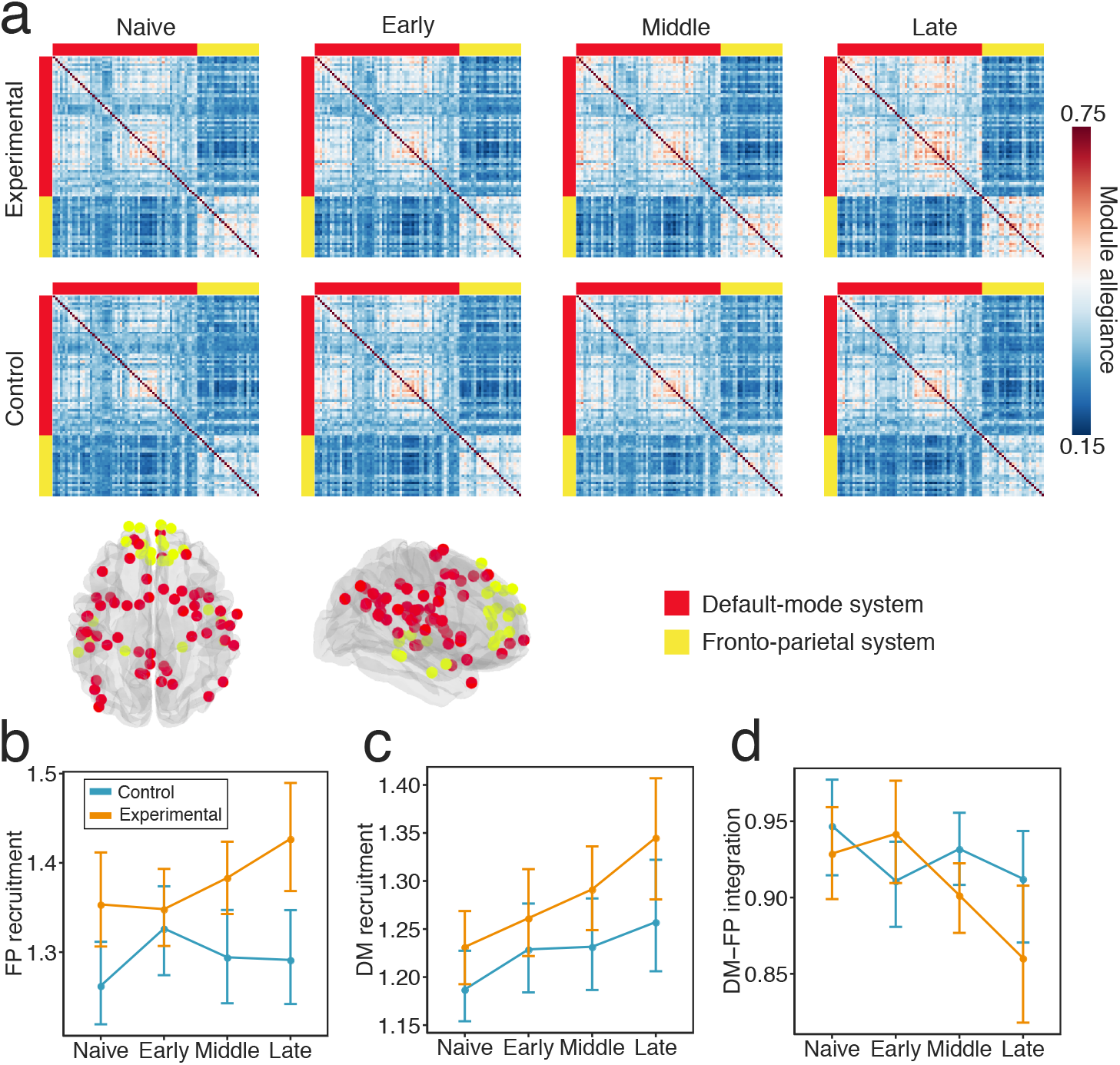
Changes in module allegiance of the fronto-parietal (FP) and default-mode (DM) systems. (**a**) Module allegiance matrices for the default mode and fronto-parietal systems. Each ij-th element of the matrix represents the probability that node *i* and node *j* are assigned to the same module within a single layer of the multilayer network. (**b**) Only the experimental group exhibited increases in fronto-parietal recruitment across sessions. (**c**) Both experimental and control groups exhibited increases in default mode recruitment between ‘Naive’ and ‘Late’ stages of training. (**d**) In both groups, the integration between the fronto-parietal and default mode systems decreased from ‘Naive’ to ‘Late’ sessions, but groups differed in the pattern of integration changes between ‘Naive’ to ‘Middle’ sessions (see also Supplementary Figure 6). Source data are provided as a Source Data file.

First, we examine dynamic topological changes in the fronto-parietal and default mode systems, which were directly related to our hypothesis. Using a multilevel model, we observed a significant session × group interaction effect when considering changes in the recruitment of the fronto-parietal system during training (*χ*^2^(3) = 9.03, *p* = 0.028; Figure 5b). The largest increase in fronto-parietal recruitment was observed in the experimental group when comparing ‘Early’ to ‘Late’ training phases (*β* = −0.07, *t*(120) = −2.892, *p* = 0.027, Bonferroni-corrected; Figure 5b). No significant changes from ‘Naive’ to ‘Late’ training phases were observed in the control group (*β* = −0.03, *t*(120) = −1.169, *p* = 1, Bonferroni-corrected). Turning to an examination of the default mode, we found a significant main effect of session (*χ*^2^(3) = 24.17, *p* < 0.0001) and of group (*χ*^2^(1) = 3.96, *p* = 0.046) on system recruitment (Figure 5c). However, the interaction effect between session and group was not significant (*χ*^2^(3) =2.66, *p* = 0.48). Planned contrasts revealed that the default mode recruitment increased steadily in both groups and we observed the largest increase between ‘Naive’ and ‘Late’ sessions (*β* = 0.09, *t*(123) 5.00, *p* < 0.0001). The experimental group displayed a higher default mode recruitment than the control group (*t*(165.6) = −3.03, *p* = 0.003). We found a significant session × group interaction effect on the integration between the frontoparietal and default mode systems (*χ*^2^(3) = 14.25, *p* = 0.0025) (Figure 5d). The integration between these two systems decreased from ‘Naive’ to ‘Late’ sessions only in the experimental group (*β*= 0.07, *t*(120) = 4.37, *p* = 0.0002, Bonferroni-corrected). However, groups differed from ‘Naive’ to ‘Early’ (*β* = 0.07, *t*(120) = 2.16, *p* = 0.03) and from ‘Early’ to ‘Middle’ sessions (*β* = −0.06, *t*(120) = −2.70, *p* = 0.02, Bonferroni-corrected): whereas the experimental group displayed an inverted U-shaped curve of integration with training, the control group displayed the opposite pattern. Collectively, these results suggest that the increase of fronto-parietal system recruitment and the decrease of integration between the default mode and fronto-parietal systems reflect training-specific changes in dual n-back task automation. In contrast, the increase in default mode system recruitment may reflect more general effects of behavioral improvement, as it was observed in both experimental and control groups.

Next, we asked whether changes in dynamic topology could be observed in other large-scale systems. Using multilevel modeling, we observed three distinct types of changes occurring over time regardless of the group (*p* < 0.05, FDR-corrected; Figure 6a-c): an increase in system recruitment, (2) an increase in the integration between task-positive systems, and (3) a decrease in the integration between default mode and task-positive systems (Supplementary Figure 7, Supplementary Table 1–2). First, we observed an increase in the recruitment beyond the default mode system – in salience, and auditory systems (Supplementary Figure 7a-c). Second, we observed an increase in the integration between task-positive systems, including fronto-parietal and salience, dorsal attention and salience, and dorsal attention and cingulo-opercular (Supplementary Figure 7d-f). Third, for the default mode system, we observed a decrease in integration with other task-positive systems: salience and cingulo-opercular (Supplementary Figure 7g-i). Additionally, we also observed a decrease in integration between the memory and somatomotor systems, and between the default mode and auditory systems (Supplementary Figure 7j-k). We observed a similar pattern of changes for the Schaefer parcellation (Supplementary Figure 17, Supplementary Figure 18a). These results suggest that the increase of within-module stability, the increase of default mode system independence from task-positive systems, and the decrease of integration between task-positive systems reflect general effects of task training.

**Figure 6:**
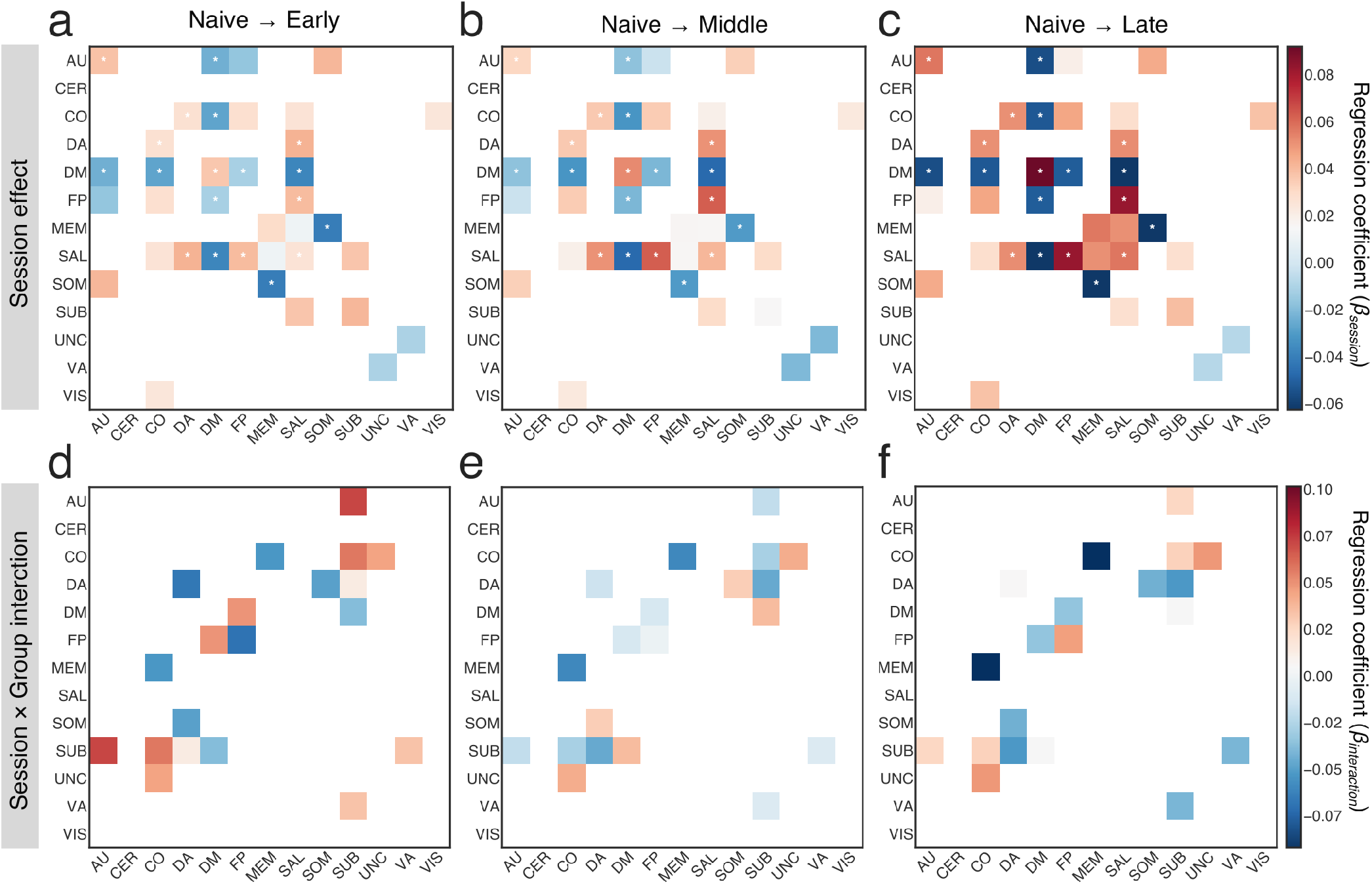
Changes of the recruitment and integration of large-scale systems. Colored tiles represent all significant effects (*p* < 0.05, uncorrected; \**p* < 0.05 FDR-corrected). (*top panel*) Here we display the significant main effects of session. Tile color codes a linear regression coefficient (*β*), for all main session effects: (**a**) from ‘Naive’ to ‘Early’, (**b**) from ‘Naive’ to ‘Middle’, and (**c**) from ‘Naive’ to ‘Late’. (*bottom panel*) Here we display the significant session × group interaction effects. Tile color codes a linear regression coefficient between groups and sessions: (**c**) from ‘Naive’ to ‘Early’, (**d**) from ‘Naive’ to ‘Middle’, and (**e**) from ‘Naive’ to ‘Late’. Abbreviations: auditory (AU), cerebellum (CER), cingulo-opercular (CO), default mode (DM), dorsal attention (DA), fronto-parietal (FP), memory (MEM), salience (SAL), somatomotor (SOM), subcortical (SUB), uncertain (UNC), ventral attention (VA), and visual (VIS). Source data are provided as a Source Data file.

We also investigated the relationship between across-session change in system recruitment or integration and across-session change in behavioral performance for all large-scale systems. For both brain and behavioral variables, we measured the change from the first (‘Naive’) to the last (‘Late’) training sessions (see Figure 7a, Supplementary Table 6). We found a significant positive correlation between change in behavior, as operationalized by a change in *d*′ (2-back minus 1-back), and change of the default mode (*r* = 0.33, *p* = 0.03, uncorrected) and salience (*r* = 0.34, *p* = 0.03; uncorrected) systems recruitment. Greater behavioral improvement was also associated with a higher increase of integration between fronto-parietal and salience systems (*r* = 0.35, *p* = 0.02, uncorrected) and a higher decrease of integration between default mode and task-positive systems: frontoparietal (*r* = −0.31, *p* = 0.04, uncorrected) and salience (*r* = −0.41, *p* = 0.006, uncorrected). Analogous relationships for default mode recruitment and default mode – fronto-parietal integration with behavioral improvement were observed for an alternative measure of performance (pRT; Supplementary Figure 9a). Note, that the correlation for the change in the *d*′ measure has opposite sign when compared to the correlation with the change of pRT, consistent with the fact that these two measures have different interpretations (the lower pRT, the better; the higher the *d*′, the better). In summary, a higher increase of stability in the default mode and salience systems, together with a decrease of default mode – task-positive systems integration may support behavioral improvement in the task, regardless of whether the task was additionally trained or not.

**Figure 7:**
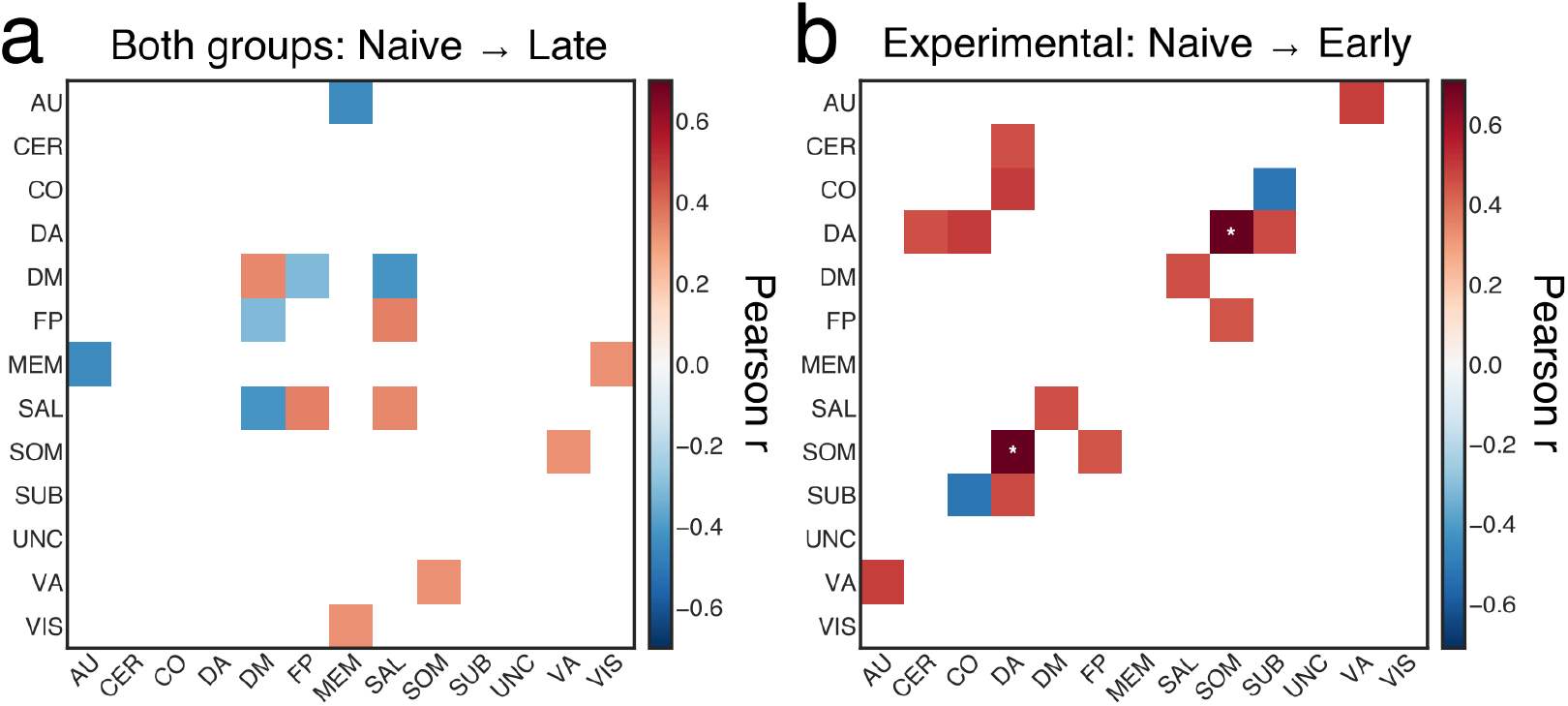
Relationship between the change in network dynamics and the change in behavior. Colored tiles represent all significant correlations (*p* < 0.05, uncorrected; \**p* < 0.05 FDR-corrected). (**a**) Pearson correlation coefficient (*r*) between the across-session changes in recruitment (or integration) and the across-session changes in *d*′ (Δ*d*′) observed for both experimental and control group. (**b**) Relationship between the changes in recruitment (or integration) and the changes in *d*′ during early phase of training of the experimental group. Abbreviations: auditory (AU), cerebellum (CER), cingulo-opercular (CO), default mode (DM), dorsal attention (DA), frontoparietal (FP), memory (MEM), salience (SAL), somatomotor (SOM), subcortical (SUB), uncertain (UNC), ventral attention (VA), and visual (VIS).Source data are provided as a Source Data file.

Finally, we also observed session × group interaction effects beyond the default mode and fronto-parietal systems (*p* < 0.05, uncorrected, Figure 6d-e, Supplementary Table 3–4). Specifically, in the experimental group, we observed a non-linear change in the integration of the subcortical system with the dorsal attention, ventral attention, cingulo-opercular, and auditory systems. An initial increase in integration with the subcortical system (from ‘Naive’ to ‘Early’) was followed by a decrease in the integration at later time intervals. Interestingly, we observed the reverse pattern for the change in integration between the subcortical and default mode systems: the integration first decreased from ‘Naive’ to ‘Early’ sessions, and then increased from ‘Early’ to ‘Middle’ sessions for the experimental group (Supplementary Figure 8; Supplementary Table 5). The pattern of changes in integration also differed between the groups, particularly so for the integration between cingulo-opercular and memory systems, cingulo-opercular and uncertain systems, and dorsal attention and somatomotor systems. These results suggest that task automation during initial stages of working memory training might also be supported by an increased communication between subcortical and other large-scale systems.

We further tested whether changes in systems recruitment or integration from ‘Naive’ to ‘Early’ sessions were associated with performance improvement displayed by the experimental group. Interestingly, we found that the behavioral change was positively correlated with change of integration between multiple systems, in particular: dorsal attention and somatomotor, dorsal attention and subcortical, fronto-parietal and somatomotor, dorsal attention and cingulo-opercular, salience and default mode. In contrast, the increase of integration of subcortical system and cingulo-opercular systems was negatively correlated with the change in task performance (Figure 7b, Supplementary Table 7). This pattern of associations between behavioral and network changes suggests that inter-systems communication might be necessary for efficient task performance during initial stages of training.

In summary, we observed two patterns of dynamic changes in network topology following working memory training. The first pattern reflects improved behavioral performance and is characterized by a gradual increase in default mode autonomy and in the integration between task-positive systems. The second pattern reflects changes related to task automation specifically in the experimental group and is characterized by non-linear changes in default mode – fronto-parietal integration, and in the integration with the subcortical system.

## DISCUSSION

In the present study, we aimed to verify the hypothesis that training on an effortful cognitive task – a dual n-back – increases the segregation of task-related functional brain networks. We examined these training-related changes utilizing both static and dynamic network approaches. While performing a dual n-back task, participants were scanned four times using fMRI: prior to training, after two weeks of training, after four weeks of training, and after six weeks of training. We examined the effect of training on whole-brain modularity, as well as on the dynamic expression of that modularity through measures of segregation and integration in large-scale systems. We found that whole-brain modularity significantly differed between task conditions, being the highest in the resting state, lower in the 1-back condition, and even lower in the 2-back condition. In the experimental group, modularity increased in response to working memory training. We also observed two patterns of changes in the dynamic network topology following training: (i) a gradual increase in the segregation of default mode and task-positive systems, and (ii) a non-linear change in the default mode - frontoparietal integration and integration of the subcortical system. The general behavioral improvement in the task in response to training was positively correlated with an increase in the recruitment of the default mode system and a decrease in its integration with the frontoparietal system. Collectively, these findings suggest that segregation of the default mode and task-positive systems supports general improvement in the task, while dynamic communication of the default mode with the fronto-parietal and subcortical systems supports more specific network changes related to automation of the working memory task.

The balance between segregated and integrated brain states is constantly re-negotiated in the face of challenges posed by the external world^11^. Together with the existence of inter-modular connections, modular organization of the brain network provides a basis for the emergence of segregated and integrated neuronal states^22^. The degree of modularity in functional brain networks can change over a variety of time scales^23^, from that of seconds as probed by intracranial recordings^24^ to that of years as driven by development^25^ or aging^26,27^. Modularity can also be modulated at an intermediate temporal scale, by task demands and cognitive effort. Several studies have reported a decrease in functional brain network modularity during increasing demands on executive function, for example by varying the level of the n-back task^2,5^. Here, we were curious to understand whether and how whole-brain network modularity changes when a demanding n-back task is intensively trained. We expected that network modularity would gradually increase during dual n-back task training, suggesting more segregated, and therefore less costly, information processing, after task automation.

We observed that modularity during the resting state was higher than during performance of the dual n-back task. Moreover, we found that the modularity during the low-demand task condition (1-back) was higher than the modularity during the high-demand task condition (2-back). Our results are consistent with previous studies providing evidence that network segregation is lowest (while integration is highest) during a demanding n-back task, when compared to a less demanding motor task or resting state^3,4^. Moreover, the observed difference between working memory loads is consistent with a previous study from^2^ who reported higher modularity during the 3-back condition compared to the 0-back condition, and also consistent with a previous study from^5^ who reported higher modularity during the 2-back condition compared to the 1-back condition. Collectively, the findings also support the Global Workspace Theory (GWT)^13^, by showing that less demanding, highly automated tasks can be performed within segregated modules, while more challenging tasks require integration between multiple modules.

Despite the consistency between our findings and prior work, it is important to note that these previous studies did not address the question of whether a fully mastered demanding cognitive task would still require a costly integrated workspace or could instead be executed within specialized brain modules. Here, our study expands upon prior work by offering the first evidence supporting the latter hypothesis. We observed that although modularity of the network generally increased through n-back training, as measured during both 1-back and 2-back conditions in the experimental group, the modularity difference between the two conditions was preserved. This finding suggests that training resulted in the increase of the baseline network segregation during the task, which supports our hypothesis that mastered cognitive tasks can be executed within a segregated network. Modularity measured during the high-demand 2-back condition after training exceeded the modularity during the low-demand 1-back condition before training. However, even if the baseline network segregation increased after the training, some level of modularity breakdown during increasing cognitive demands seems to be induced.

Importantly, modularity is also altered in patients with disorders of mental health or patients sustaining brain injury. Studies have found that modular organization of a network is disrupted in patients with cognitive control deficits^28^, and increases over the early stage of stroke recovery in a manner that is related to the recovery of higher cognitive functions^29^. Further longitudinal studies in these patient populations could provide clarity on the role of modularity – and its variation over a range of time scales – in higher-order cognitive function. Our findings suggest that there is a possibility to increase brain network modularity via intensive working memory training. This phenomenon may have potential beneficial implications for designing cognitive training interventions to prevent aging-related cognitive decline, reduce cognitive control deficits, or intensify effects of neurorehabilitation through increasing brain plasticity. To verify this conjecture, future studies should examine the direct effect of training-induced increase of brain modularity in healthy ageing and clinical populations.

Interestingly, we did not observe differences between the experimental and control group in the increase of network modularity. The control group displayed a small increase of modularity in the 1-back condition, suggesting that the segregation of the functional brain network may increase rapidly, also in response to repeated exposure to the task. The control group performed the dual n-back task four times during scanning sessions, which resulted in a small behavioral improvement. This result suggests that the increase of network segregation may be sensitive to varying intensity of training in the task. Future studies with a larger sample size should examine whether such gradation exists.

The modular structure of functional brain networks is not static, but instead undergoes dynamic reconfiguration throughout a range of cognitive processes^1,30–34^. Recently developed dynamical approaches to study brain networks are sensitive to the temporal nature of the underlying neural signal, and therefore can be used to probe the fluctuating patterns of connectivity elicited by task performance. Using just such a dynamical approach, Bassett et al.^7^ showed that the modular structure of human brain functional networks fluctuates appreciably during motor-visual learning, and moreover that the degree of fluctuations changes during a 6-week training paradigm. Task-relevant, motor and visual networks exhibited increasing autonomy as the duration of training increased, marking the emergence of automatic behavioral responses. In light of this prior work, we hypothesized that networks relevant to working memory function – including the fronto-parietal and default mode systems – would increase their autonomy after extensive training on a working memory task.

**Figure 8:**
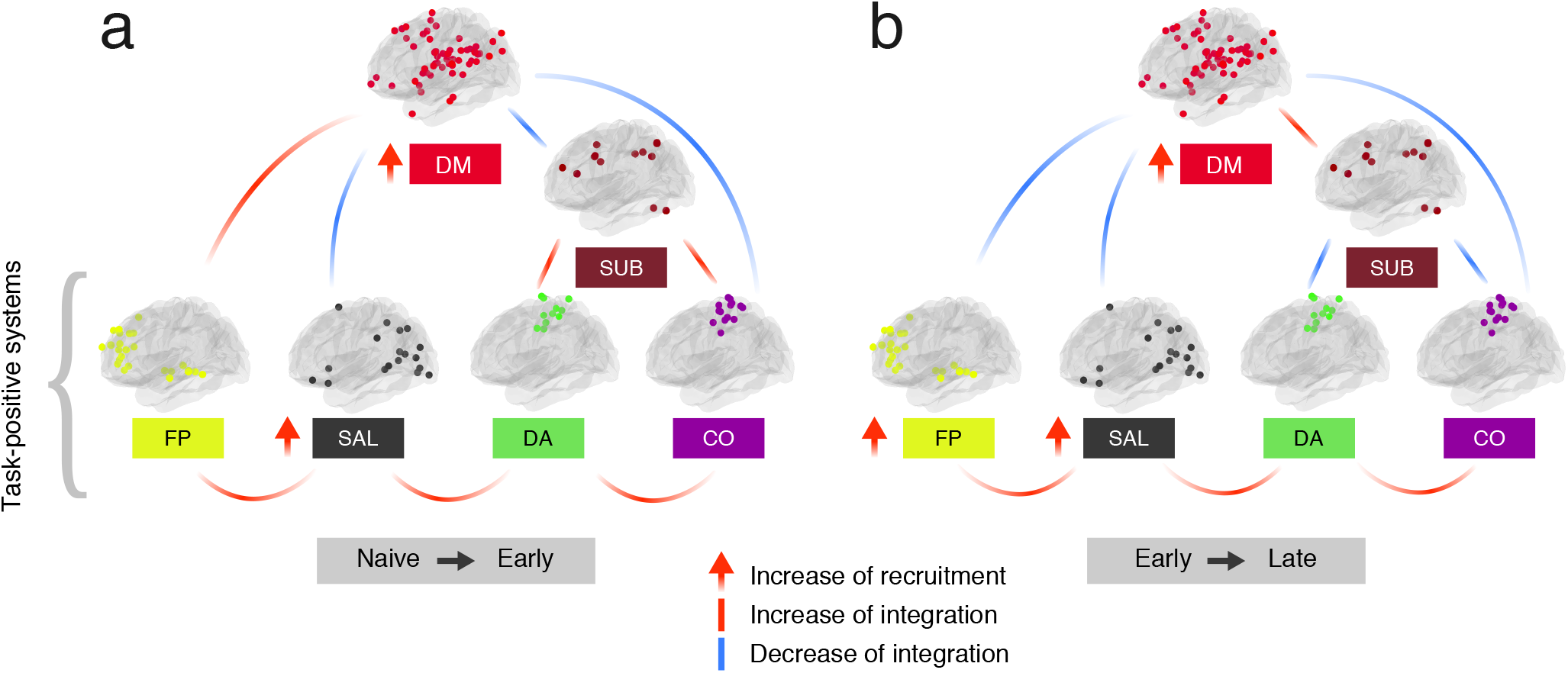
Schematic diagram summarizing the main changes in recruitment and integration of large-scale systems observed over the course of working memory training. We observed a gradual increase of the integration between task-positive systems (fronto-parietal - FP, salience - SAL, dorsal attention - DA, and cingulo-opercular - CO), greater recruitment of the default mode (DM) system, and decreased DM-CO and DM-SAL integration. In the early phase of training (**a**) the experimental group displayed an increased FP-DM integration, and increased integration of the subcortical (SUB) system with the DA and CO systems. In the late stage of training (**b**), the FP system reduced its integration with the DM system and the SUB system increased its integration with the DM system, while decreasing coupling with task-positive systems.

Here, we used a multilayer community detection algorithm to determine whether modular structure of large-scale systems change in response to n-back training. We further applied multilevel modeling^35^ to test for possible group and session differences in the dynamic network measures while controlling for differences in individual baseline values. In testing our hypothesis, we held in mind the observations of previous studies, which have noted that the fronto-parietal and default mode systems can both cooperate and compete during tasks that require cognitive control, such as the n-back task^15,36^. Understanding the nature of interactions between these two systems is therefore essential for explaining the neural adaptation that occurs in response to evolving cognitive demands. It is also not known whether dynamic interactions between these two networks may evolve during cognitive training. Using dynamic network metrics, we showed that the default mode system increased its recruitment in both groups, indicating that regions within this system were coupled with other communities less often. Moreover, the experimental group displayed an increased fronto-parietal recruitment and an inverted U-shaped curve of integration between the default mode and fronto-parietal systems with training. Enhanced default mode intra-communication and decreased intercommunication with the fronto-parietal system were associated with better behavioral outcomes after training. We also observed significant changes in dynamic network topology beyond the fronto-parietal and default mode systems. In particular, regardless of the group, we observed an increased recruitment of the salience and auditory systems, decreased integration between the default mode and other task-positive systems (including salience and cingulo-opercular), and increased integration between task-positive systems (including frontoparietal and salience, dorsal attention and salience, dorsal attention and cingulo-opercular). These results suggest the existence of the trade-off between segregation and integration: whereas segregation increases between some systems, the integration increases or decreases between others.

Some studies suggest that competitive interactions between the task-positive fronto-parietal system and the task-negative default mode system might be essential for higher order cognitive functions^15,36^. The frontoparietal system is composed of spatially distributed brain areas including the lateral prefrontal cortex, anterior cingulate, and inferior parietal cortex^37^. Its activity is commonly linked to the performance of tasks requiring cognitive control, such as the n-back working memory task^37,38^. Prior work offers evidence that the fronto-parietal system is highly flexible and dynamically interacts with other systems in response to the changing demands of cognitive tasks^1,31^. In contrast, the default mode system exhibits high activity during internally directed cognition, such as mind wandering and autobiographical memory^39^. The default mode system is composed of spatially distributed brain areas including the medial prefrontal cortex, posterior cingulate, lateral parietal cortex, and both lateral and medial temporal cortices^39,40^. The default mode’s activity is frequently anticorrelated with the activity of systems that engage in demanding cognitive tasks such as the fronto-parietal and dorsal attention systems^41^. Recent studies, however, challenge a common view about existing antagonism between default mode and frontoparietal systems, suggesting that the interaction between these two systems is necessary for efficient behavioral control^42,43^. Recent findings confirm that default mode regions display a positive coupling with task-positive brain systems during working memory task performance^44,45^ and may dynamically switch their connections to support inter-module communication in high-demand n-back task conditions^5,46^. By using dynamic network approach, we were able to track both task-related and training-related fluctuations of modular network structure during the working memory task performance. Our observations expand upon prior studies by demonstrating that the increase in default mode segregation and decrease of integration between the default mode and fronto-parietal systems may be an indicator of behavioral improvement during working memory training. Moreover, the previous study reported the relationship between the default mode connectivity changes and static modularity changes during n-back task^5^. In our exploratory analysis, we also showed that default mode recruitment fluctuated between task conditions and was significantly higher in the 1-back condition than in the 2-back condition (Supplementary Figure 19) and, similar to modularity, increased steadily in both groups. Here we also observed a positive relationship between the change in default mode recruitment and change of modularity from ‘Naive’ to ‘Late’ session (Supplementary Figure 21). As we did not observe the relationship between changes of modularity and behavioral improvement, we may conclude that studying the dynamics of modular network structure enables a better prediction of behavioral outcomes in response to training.

Our results are also consistent with prior observations that the default mode and fronto-parietal systems may interact in a task-dependent manner with the salience, cingulo-opercular, and dorsal attention systems^15,47^. Bressler and Menon^47^ proposed a model whereby efficient cognitive control is supported by the dynamic switching between functionally segregated fronto-parietal and default mode systems mediated by cingulo-opercular and salience systems. Cocchi et al.^15^ proposed that task-related reconfiguration is possible through flexible interactions within and between overlapping meta-systems: (i) the executive meta-systems, responsible for the processing of sensory information, and (ii) the integrative meta-system, responsible for flexible integration of brain systems. These two metasystems are composed of transient coupling between three large-scale systems: the frontoparietal system, the cingulo-opercular/salience system, and the default mode system. During high-demand task conditions the executive meta-system is formed by extensive interactions between fronto-parietal and cingulo-opercular/salience systems, and the default mode system is more segregated and less integrated with the fronto-parietal system^48^. Our results extend these findings by presenting the evolving reconfigurations of large-scale networks during mastery of the working memory task. We showed that regardless of the group the default mode system reduced coupling with the cingulo-opercular and salience systems. These results suggest that increased segregation of the default mode and task-positive networks may be a consequence of more efficient task performance. A similar pattern of changes was observed across two different subdivisions of the cortex into systems (Power and Schaefer), together suggesting that the salience and cingulo-opercular systems that are thought to be responsible for switching between antagonistic frontoparietal and default mode systems, appear to be more integrated with the fronto-parietal system and less integrated with the default mode system. This pattern of relations may be due to diminished requirements for switching between these two systems when the task is well learned.

Similar to modularity, the lack of group differences in the pattern of these changes suggests that the increased DM autonomy and increased integration of task-positive systems might be related to a general improvement in task performance. Such behavioral improvement, although much smaller than in the experimental group, was also observed in the control group during the 2-back condition. As participants performed the task four times in the scanner, they inevitably trained the task to a small extent. The presence of network reorganization in the control group may suggest that changes in DM autonomy and integration of task-positive systems occur relatively fast, even when the training is not intense. As participants of our study were scanned in 2-week intervals, we could not capture what behavioral improvement is necessary to invoke such network reorganization. To better understand the dynamics of these neuroplastic changes, future studies should examine day-to-day network reorganization in response to training with different intensities.

We also observed that groups differed in patterns of changes in the subcortical system coupling. Specifically, the experimental group displayed an inverted U-shaped curve of changes in (i) the integration between the subcortical system and the dorsal attention system, and (ii) the integration between the ventral attention system and the cingulo-opercular system; notably, the control group displayed the opposite pattern. We observed the opposite effect for coupling between the subcortical and default mode systems. Non-linear changes in subcortical activity were also observed in previous studies of the effects of working memory training^49^. Consistent with our results, Kühn et al.^49^ found that activity in the subcortical regions increased after one week of working memory training and decreased after 50 days of training. Previous studies suggested that subcortical activity can mediate changes in working memory ability^50^. Because an inverted U-shaped curve of changes in fronto-parietal activity was also observed following working memory training^49,51^, we speculate that subcortical activity may influence changes in the fronto-parietal system. Yet, results based on observation of brain activity changes cannot provide information on how these two systems interact. Our results show that in the initial training phase, the subcortical system switched coupling from task-positive systems to the default mode system. We observed the opposite pattern for the fronto-parietal system, which instead first increased and then decreased its interaction with the default mode system. We speculate that the subcortical system supports segregation of the task-positive and default mode systems. Future studies using effective connectivity approach could examine whether such a cause and effect relationship exists.

The dynamic network approach extends our understanding of training-related changes in brain function. Studies focusing on changes in brain activity during a working memory training reported a decrease of task-positive systems activation^49,51^, commonly interpreted as a reflection of increased neural efficiency within systems engaged in the task^52^. Here, we reported a similar effect using a standard GLM-based approach (see Supplementary Figure 10–11, Supplementary Table 8–9). However, we also showed that our findings on the dynamic network changes can not be simply explained by the changes in brain activity (Supplementary Figure 12). The fronto-parietal system dynamically interacts with other large-scale systems^15^, and it is reasonable to expect that working memory training might influence interactions in the whole network. We observed training-related increases in the segregation of the default mode and task-positive systems that suggest more efficient and less costly processing within these systems after training. Accordingly, greater segregation of the default mode system and task-positive systems and smaller integration between these systems were associated with behavioral performance improvement. Moreover, we showed that an increase of integration between multiple large-scale systems in early phase of training was related to a greater behavioral improvement in the experimental group, indicating that some level of network integration is necessary when the task is not fully automated. Taken together, the dynamic network approach provides a unique insight into the plasticity and dynamics of the human brain network.

## METHODS

### Subjects

Fifty-three healthy volunteers (26 female; mean age: 21.17; age range: 18–28 years) were recruited from the local community through word-of-mouth and social networks. All participants were right-handed, had normal or corrected-to-normal vision, and had no hearing deficits. Seven participants did not complete the study: one due to brain structure abnormalities detected at the first scanning session, and six due to not completing the training procedure. The final sample consisted of forty-six participants who completed the entire training procedure, participated in all four fMRI scanning sessions, and had no history of neurological or psychiatric disorders nor gross brain structure abnormalities. After the first fMRI scan, participants were matched by sex and randomly assigned to one of the two training groups: experimental and control (see next section on **Experimental Procedures**). Each group consisted of 23 subjects with no group differences in age (two-sample t-test: *t*(42.839) = 0.22, *p* = 0.83) or fluid intelligence (two-sample t-test: *t*(42.882) = 0.51, *p* = 0.61) as measured by Raven’s Advanced Progressive Matrices (RAPM)^53^. Informed consent was obtained in writing from each participant, and ethical approval for the study was obtained from the Ethics Committee of the Nicolaus Copernicus University Ludwik Rydygier Collegium Medicum in Bydgoszcz, Poland, in accordance with the Declaration of Helsinki.

### Experimental Procedures

The study was performed at the Centre for Modern Interdisciplinary Technologies, Nicolaus Copernicus University in Toruń (Poland). Each participant who completed the entire study procedure attended a total of 24 meetings at the laboratory. During the first meeting, participants were familiarized with the study procedure and timeline, and were asked to provide basic demographic information and informed consent. During the second meeting, participants performed fluid intelligence testing with RAMP^53^. Then, participants were scheduled for fMRI testing, which was performed before training, after two weeks of training, after four weeks of training, and after 6 weeks of training. Each fMRI session was scheduled to be on the same day of the week and at the same hour for each participant. These schedules varied in exceptional cases (holidays, illness of participant, emergency). However, scanning procedures were always performed between 24h to 48h after the last training session. After the first fMRI session, participants were randomly assigned to one of two training groups: (1) experimental, which trained working memory with an adaptive dual n-back task^14^, and (2) a passive control group which interchangeably performed an auditory and spatial 1-back task. We included this second group to control for differences in the effect of training on task performance and fMRI signatures driven by repeated exposure to a task.

Two versions of the dual n-back task were used: (1) an adaptive dual n-back was used in the training sessions of the experimental group only, and (2) an identical dual n-back task with two conditions (1-back and 2-back) used during fMRI scanning of both groups. Both scanning and training versions of the dual n-back task consisted of visuospatial and auditory tasks performed simultaneously. Visuospatial stimuli consisted of 8 blue squares presented sequentially for 500 ms on the 3 × 3 grid with a white fixation cross in the middle of the black screen; auditory stimuli consisted of 8 Polish consonants (b, k, w, s, r, g, n, z) played sequentially in headphones. Participants were asked to indicate by pressing a button with their left index finger whether the letter heard through the headphones was the same as the letter n-back in the sequence, and by pressing a button with their right index finger to indicate whether the square on the screen was in the same location as the square n-back in the sequence.

In the training version of the task the *n*-level of the dual n-back task increased adaptively when participants achieved 80% correct responses in the trial, and the *n* level decreased when participants made more than 50% errors in the trial. After each trial, the *n* level achieved by a participant was recorded, and the mean *n* level during each of 18 training session was used later to calculate the total training progress (Supplementary Figure 1, Supplementary Figure 2). Participants from the control group performed a single 1-back with auditory or visuospatial stimuli variants. To minimize boredom of participants, the order of the 1-back variants was randomly selected at the beginning of each training session. Therefore, each participant from the control group had the same number of training trials on single auditory and visuospatial n-back tasks. Participants completed a total of 18 sessions (30 min each) under the supervision of the experimenter. Each participant completed 20 blocks (each consisting of 20 + *n* trials, depending on the *n* level achieved by the participant) of the n-back task during each training session. The study was double-blind; the experimenter performing the fMRI examination was not aware of the group assignment of the participants, and participants were not aware that the study was designed in a way that there were two groups (experimental and control). The apparatus used in the study consisted of two 17” Dell Inspiron Laptops, and two pairs of Sennheiser headphones. Stimulus delivery was controlled by a Python adaptation of the dual n-back task used by Jaeggi et al.^14^ (http://brainworkshop.sourceforge.net/).

In the fMRI scanning version of the task, participants performed the dual n-back task with two levels of difficulty: 1-back and 2-back. Each session of the task consisted of 20 blocks (30 s per block; 12 trials with 25% of targets) of alternating 1- and 2-back conditions. To enable for dynamic network comparison across blocks, we did not add any systematic variation to block length and block order. The instruction screen was displayed for 4,000 ms before each block, informing the participant of the upcoming condition. Both visual and auditory stimuli were presented in a pseudo-random order. Participants were asked to push the button with their right thumb if the currently presented square was in the same location as the previous square (1-back) or two squares back in the sequence (2-back) and, at the same time, push the button with their left thumb when the currently played consonant was the same as the previous consonant (1-back) or two consonants back (2-back). The participants had 2,000 ms to respond, and were instructed to respond as quickly and accurately as possible. The experimental protocol execution and control (stimulus delivery and response registration) employed version 17.2. of Presentation software (Neurobehavioral Systems, Albany, NY), as well as MRI compatible goggles (visual stimulation), headphones (auditory stimulation), and response grips (response registration) (NordicNeuroLab, Bergen, Norway). Before each scanning session, participants performed a short dual n-back training session outside the fMRI scanner to (re-)familiarize them with the rules of the task.

All participants received equal monetary remuneration (200 PLN) for study participation together with a radiological description and a CD containing their anatomical brain scans.

### Data acquisition

Neuroimaging data were collected using a GE Discovery MR750 3 Tesla MRI scanner (General Electric Healthcare) with a standard 8-channel head coil. Structural images were collected using a three-dimensional high resolution T1-weighted gradient-echo (FSPGR BRAVO) sequence (TR = 8.2 s, TE = 3.2 ms, FOV = 256 mm, flip angle = 12 degrees, matrix size 256 × 256, voxel size = 1 × 1 × 1 mm, 206 axial oblique slices). Functional scans were obtained using a T2*-weighted gradient-echo, echo-planar imaging (EPI) sequence sensitive to BOLD contrast (TR = 2,000 ms, TE = 30 ms, FOV = 192 mm, flip angle = 90 degrees, matrix size = 64 × 64, voxel size 3 × 3 × 3 mm, 0.5 mm gap). For each functional run, 42 axial oblique slices were acquired in an interleaved acquisition scheme, and 5 dummy scans (10 s) were obtained to stabilize magnetization at the beginning of the EPI sequence. Resting state (10 min 10 s, 305 volumes) data was acquired at the beginning of each scanning session. During the resting state, participants were asked to focus their eyes on the fixation cross in the middle of the screen. The dual n-back task data (11 min 30 s; 340 volumes) were acquired using the same data acquisition settings (see Experimental Procedures for the task description.

### Behavioral performance

To measure behavioral performance in the dual n-back scanning sessions, we incorporated *d*’, a signal detection theory statistic^16^. This measure combines both response sensitivity and response bias. For every subject, session, task condition, and stimulus modality, we first divided all responses into four categories: (1) hits – button press for targets, (2) misses – lack of response for targets, (3) false alarms – button press for non-targets and (4) correct rejections – lack of response for nontargets. We defined hit rate *H* and false alarm rate *F* as:

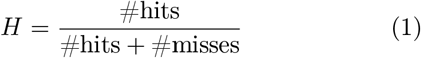

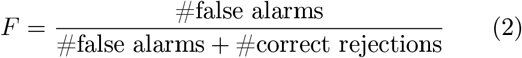

We calculated *d′* measure as:

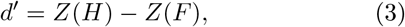

where *Z*(*x*) is the inverse of the cumulative Gaussian distribution. To get finite values of *d′* for the situations in which *H* or *F* was equal to 0 or 1, we used modified values of either 0.01 or 0.99 instead. For each participant, we calculated average *d′* for both modalities to represent a cumulative measure of performance during the dual n-back task. We also calculated behavioral performance using an alternative measure, penalized reaction time (pRT), which incorporates a measure of accuracy (see Supplementary Figure 4 for variability changes for these measures).

### Data processing

After converting from DICOM to NifTI format, functional and anatomical data were structured according to the BIDS (Brain Imaging Data Structure) standard^54^ and validated with BIDS Validator (https://bids-standard.github.io/bids-validator/). Neuroimaging data was preprocessed using fMRIPrep 1.1.1^55^ a Nipype^56^ based tool. See Supplementary Methods for details on anatomical data processing. Functional data was slice time corrected using 3dTshift from AFNI v16.2.07^57^ and motion corrected using MCFLIRT (FSL v5.0.9,^58^. This process was followed by co-registration to the corresponding T1w using boundary-based registration^59^ with 9 degrees of freedom, using bbregister (FreeSurfer v6.0.1). Motion correcting transformations, BOLD-to-T1w transformation and T1w-to-template (MNI) warp were concatenated and applied in a single step using antsApplyTransforms (ANTs v2.1.0) employing Lanczos interpolation.

Physiological noise regressors were extracted by applying CompCor^60^. Principal components were estimated for the two CompCor variants: temporal (tCom-pCor) and anatomical (aCompCor). A mask to exclude signal with cortical origin was obtained by eroding the brain mask, ensuring that it only contained subcortical structures. Six tCompCor components were then calculated including only the top 5% variable voxels within that subcortical mask. For aCompCor, six components were calculated within the intersection of the subcortical mask and the union of the CSF and WM masks calculated in T1w space, after their projection to the native space of each functional run. Frame-wise displacement^61^ (FD) was calculated for each functional run using the implementation of Nipype. The internal operations of fMRIPrep use Nilearn^62^, principally within the BOLD-processing workflow. For more details of the pipeline see https://fmriprep.readthedocs.io/en/latest/workflows.html.

Non-smoothed functional images were denoised using Nilearn^62^ and Nistats. We implemented voxelwise confound regression by regressing out (1) signals from six aCompCor components, (2) 24 motion parameters representing 3 translation and 3 rotation timecourses, their temporal derivatives, and quadratic terms of both, (3) outlier frames with FD >0.5mm and DVARS (Derivative of rms VARiance over voxelS)^63^ with a threshold of ±3 SD, together with their temporal derivatives, (4) task effects and their temporal derivatives^64^, and (5) any general linear trend. Time-series were filtered using 0.008-0.25 Hz band-pass filter. We excluded four high motion participants (2 from the control group, and 2 from the experimental group) with a mean FD larger than 0.2 mm and more than 10%of outlier volumes in any scanning session (Supplementary Figure 5).

### Functional connectivity estimation

Functional connectivity is a measure of the statistical relation between time-series of spatially distinct brain regions. Time-series can be defined as signals from single voxels or as the mean of the signals from anatomically or functionally defined groups of voxels, also known as brain parcels^65^. Here, we used a functional brain parcellation composed of 264 regions of interests (ROIs) provided by Power et al.^19^. This parcellation was based on meta-analysis and has previously been used in many studies focused on task-based network reorganization^2,5,31^. To validate our results, we also used a 300-ROI parcellation provided by Schaefer et al.^66^, which is based on transitions of functional connectivity patterns.

We created *N* × *N* correlation matrices by calculating the Pearson’s correlation coefficient between the mean signal time-course of region *i* and the mean signal time-course of region *j*, for all pairs of ROIs (*i,j*). We retained only positive correlations for further analysis. In the case of the dual n-back task, we employed a weighted correlation measure, to control for delays due to the hemodynamic response function (HRF)^64^. In this procedure, we first convolved task block regressors with the HRF and applied a filter to retain only positive values of the resultant time-series. Then, original timeseries were filtered according to the task condition and positive values of the HRF-convolved time-series. Next, the weighted correlation coefficient was calculated between the concatenated block time series, with weights taken from the corresponding HRF-convolved signals. Finally, Fisher’s transformation was employed to convert Pearson’s correlation coefficients to normally distributed *z*-scores. This procedure resulted in 264 × 264 (Power parcellation) and 300 × 300 (Schaefer parcellation) correlation matrices for each subject, session, and task condition (resting-state, 1-back, 2-back). For the dynamic network analyses, we calculated the weighted correlations for each block of the n-back task, resulting in 264 × 264 × 20 and 300 × 300 × 20 matrices, where the third dimension represents the number of task blocks (20 interleaved blocks of 1-back and 2-back).

### Static modularity

To calculate the extent of whole-brain network segregation, we employed a Louvain-like community detection algorithm^67^ to optimize a common modularity quality function^17^. This algorithm partitions the network into communities, where nodes in a given community are highly interconnected among themselves, and sparsely interconnected to the rest of the network. The modularity quality index, *Q*, to be optimized was defined as follows:

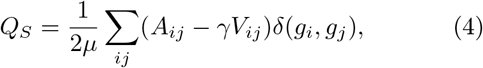

where 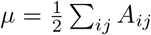 is the total edge weight of the network, *A_ij_* is the strength of the edge between node *i* and node *j*, and *γ* is the structural resolution parameter. The Kronecker delta function *δ*(*g_i_*, *g_j_*) equals one if nodes *i* and *j* belong to the same module, and equals zero otherwise. The term *V_ij_* represents the connectivity strength expected by chance in the configuration null model:

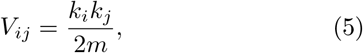

where *k_i_* and *k_j_* are the weighted degrees of nodes *i* and *j*, respectively, and 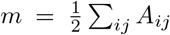 is the sum of all nodal weighted degrees.

Since the Louvain algorithm is non-deterministic, we run it 100 times, and then consider the network partition with the highest modularity score across these runs. It is important to note that the values of graph theoretical metrics can vary markedly depending on the sum of connection strengths in the network^68^. To take this effect into account, we normalized each individual modularity value against a set of modularity values calculated for randomly rewired networks^18^. For this purpose, we created 100 null networks using random rewiring of each original functional network. Then, modularity scores were calculated for each null network, thereby creating a null distribution. Finally, we normalized modularity values by dividing them by the mean of the corresponding null distribution.

### Multilayer modularity

To calculate measures of recruitment and integration, we performed multilayer modularity maximization used a generalized Louvain-like community detection algorithm introduced by^20^. This algorithm allows the optimization of a modularity quality function on a network with multiple layers. In our study, networks calculated for each separate block were considered as consecutive layers of the multilayer network. For each subject, session, and multilayer network, we ran 100 optimizations of the modularity quality function, defined as:

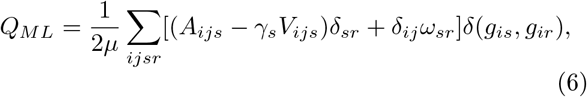

where *A_ijs_* represents the element of the adjacency matrix at slice *s*, *V_ijs_* represents the element of the null model matrix at slice *s*, *g_ir_* provides the community assignment of node *i* in slice *r*, 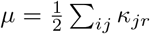 is the total edge weight of the network, where *κ_js_* = *k_js_* + *c_js_* is the strength of node *j* in slice *s*, the *k_js_* is the interslice strength of node *j* in slice *s*, and 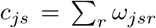. For all slices we used the Newman-Girvan null model, also known as the configuration model, defined as:

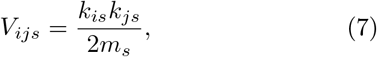

where 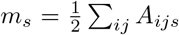 is the total edge weight of slice *s*. In this optimization, there are two free-parameters: *γ_s_* and *ω_jsr_*. The parameter *γ_s_* is the structural resolution parameter for slice *s*, and the parameter *ω_isr_* represents the connection strength between node *j* in slice *s* and node *j* in slice *r*. These two parameters can be used to tune the size of communities within each layer and the number of communities detected across all layers, respectively. Here, in line with previous studies we set *γ* = 1^21^. Due to the interleaved nature of our experimental design, *ω* = 1 for slices from the same task condition, and *ω* = 0.5 for slices from different task conditions.

### Network diagnostics

Multilayer community detection results in a single module assignment *N* × *T* matrix, where each matrix element represents the module assignment of a given node for a given slice. To summarize the dynamics of module assignments for each subject and session, we calculated an *N* × *N* module allegiance matrix, *P*, where the element *P_ij_* represents the fraction of network layers for which node *i* and node *j* belong to the same community^7,21^:

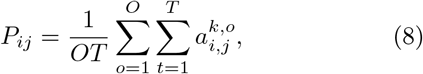

where *O* is the number of repetitions of the multilayer community detection algorithm (here, *O* = 100), and T is the number of slices (here 20 task blocks). For each optimization *o* and slice *t*,

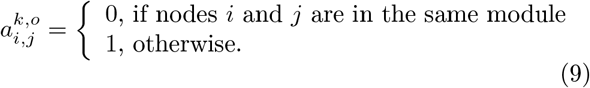

To characterize the dynamics of large-scale systems recruitment and integration, we employed methods of functional cartography^7,21^. These measures allow us to summarize how often regions from the system of interest are assigned to the same module. We can define the recruitment of system *S* as:

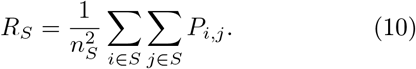

The recruitment of system *S* is high when regions within the system tend to be assigned to the same module throughout all task blocks. Similarly, we can define the integration coefficient between system *S_k_* and system *S_l_*

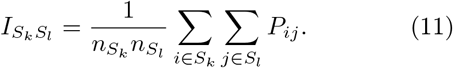

Systems of interest are highly integrated when regions belonging to two different systems are frequently assigned to the same community.

To remove potential bias introduced by the differences in the number of nodes within each system, we used permutation approach to normalize values of recruitment and integration coefficients. For each subject and session, we created *N_perm_* = 1000 null module allegiance matrices by randomly permuting correspondence between ROIs and large-scale systems. We then calculated functional cartography measures for all permuted matrices. This procedure yielded null distributions of recruitment and integration coefficients resulting solely from the size of each system. In order to obtain normalized values of *R_s_* and *I_S_k_S_t__* we divided them by the mean of the corresponding null distribution. We also calculated recruitment and integration coefficients based on multilevel community detection for signed networks (Supplementary Methods, Supplementary Figure 13-15).

### Statistical modeling

Due to the nested nature of the study data, we used two-level (trials nested within participants) and three-level (trials nested within sessions nested within participants) multilevel models^35^ (MLM) at four points during our analysis of the data. In all cases, random intercepts were estimated. The significance of models was estimated with chi-square tests, where models with increasing complexity were compared and the resulting value of Likelihood Ratio Test (*χ*^2^) and corresponding *p*-value were reported^69^. The MLM analysis was performed using *nlme*^70^ R package.

#### Behavioral changes during training

To investigate behavioral changes in behavioral performance depending on the session, task condition, and group, we used a three-level multilevel model with *d′* as the dependent variable and with group (2 factors: experimental and control), condition (2 factors: 1-back and 2-back, reference category: 1-back), and session (4 factors: Naive, Early Middle, Late; reference category: Naive) as independent variables. In addition to the main effects (group, condition, session), we included the following interaction terms: group × session, condition × session, group × condition, and group × condition × session.

#### Modularity at baseline

To investigate the dependence of static modularity at baseline on task condition, we used a two-level multilevel model with static modularity as the dependent variable and with task condition (3 factors: rest, 1-back, 2-back, two orthogonal contrasts: rest vs. 1-back and 2-back, 1-back vs. 2-back) as the independent variable. The main effect of condition was tested.

#### Training-dependent changes in static modularity

To investigate the dependence of static modularity on the session, task condition, and group, we used a three-level multilevel model with static modularity as the dependent variable and with group (2 factors: experimental and control), condition (2 factors: 1-back and 2-back), and session (4 factors: Naive, Early Middle, Late, reference category: Naive) as independent variables. In addition to the main effects (group, condition, session), we included the following interaction terms: group × session, condition × session, group × condition, and group × condition × session.

#### Changes in dynamic network metrics

To investigate changes in the integration and recruitment of large scale systems, we used a two-level multilevel model with the diagnostic measure (recruitment or integration) as the dependent variable and with group (2 factors: experimental and control) and session (4 factors: Naive, Early Middle, Late, reference category: Naive) as independent variables. In addition to the main effects (group, session), we included the following interaction term: group × session.

### Data availability

The raw behavioral data and fMRI results are available for download at https://osf.io/wf85u/ (DOI 10.17605/OSF.IO/WF85U). The source data underlying Figures 2-7, and Supplementary Figures 1-21 are provided as a Source Data file. The raw fMRI data are available from the corresponding author on request.

### Code availability

All code used for neuroimaging and behavioral data processing and statistical data analyses are publicly available at https://osf.io/wf85u/ (DOI 10.17605/OSF.IO/WF85U).

## ACKNOWLEDGMENTS

The study was supported by the National Science Centre, Poland (2015/17/N/HS6/03549). K.F. was supported by National Science Centre, Poland (2015/17/N/HS6/03549, 2017/24/T/HS6/00105) and Foundation for Polish Science, Poland (START 23.2018). Calculations have been carried out using resources provided by Wroclaw Centre for Networking and Supercomputing (http://wcss.pl), grant No. 467. D.S.B. also acknowledges support from the John D. and Catherine T. MacArthur Foundation, the Alfred P. Sloan Foundation, the ISI Foundation, the Paul Allen Foundation, the Army Research Laboratory (W911NF-10-2-0022), the Army Research Office (Bassett-W911NF-14-1-0679, Grafton-W911NF-16-1-0474, DCIST-W911NF-17-2-0181), the Office of Naval Research, the National Institute of Mental Health (2-R01-DC-009209-11, R01-MH112847, R01-MH107235, R21-M MH-106799), the National Institute of Child Health and Human Development (1R01HD086888-01), the National Institute of Neurological Disorders and Stroke (R01 NS099348), and the National Science Foundation (BCS-1441502, BCS-1430087, NSF PHY-1554488 and BCS-1631550). We also thank Maja Dobija, Alex Lubiński, Stanisław Narębski, Monika Muchlado, Bożena Pięta, and Adrianna Przybysz for their assistance in conducting training sessions, and Jaromir Patyk for technical support. The content is solely the responsibility of the authors and does not necessarily represent the official views of any of the funding agencies.

## AUTHOR CONTRIBUTIONS

K.F. provided funding for the study. K.F. designed the study with the contribution from S.K. K.F. and K.B. collected and processed data. K.F. performed network and statistical analyses with the support and contributions from D.S.B., X.H., D.M.L., and K.B. K.F wrote the manuscript. D.S.B., W.D., K.B., X.H., D.M.L, and S.K. revised the manuscript.

## COMPETING INTERESTS

The authors declare no competing interests.

## SUPPLEMENTARY INFORMATION

### 1 Supplementary Figures

**Supplementary Figure 1:**
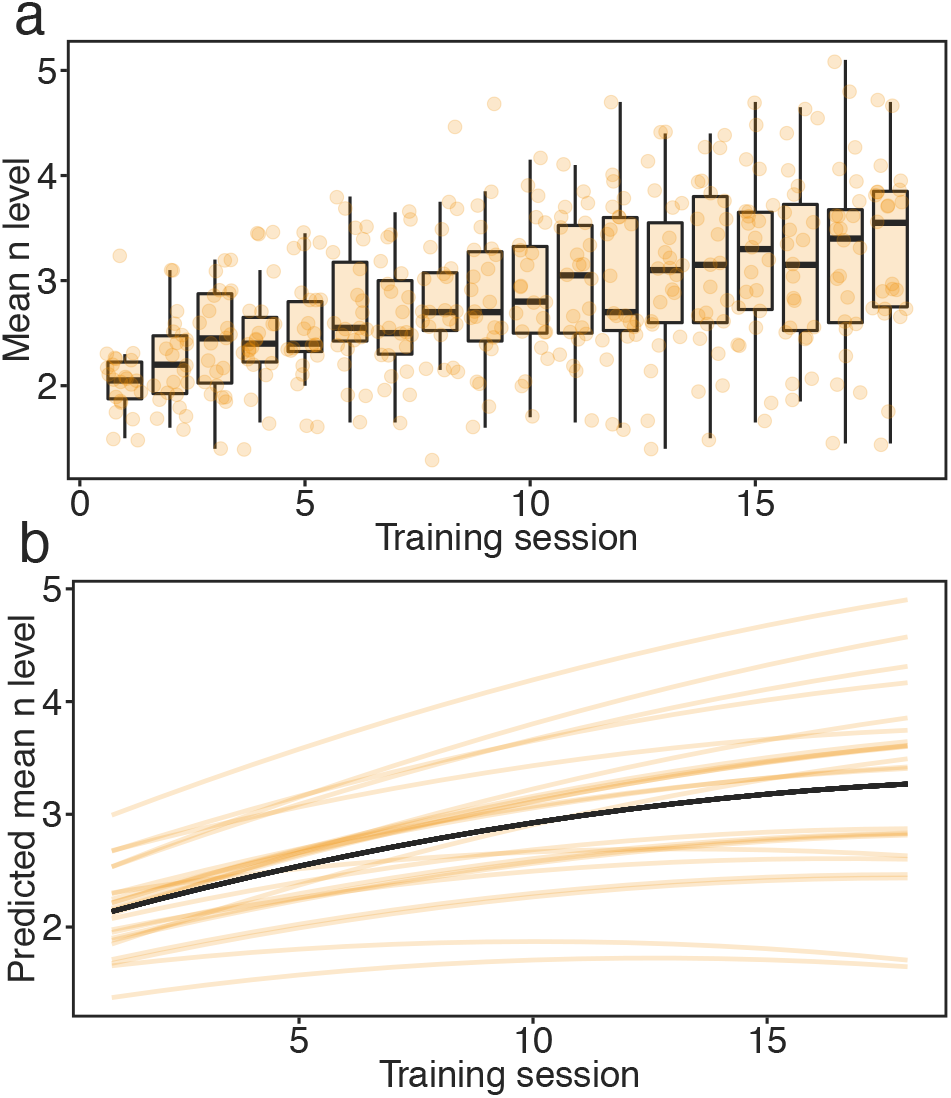
Behavioral performance during dual n-back training. The performance was measured as a mean n-back level achieved during each trial of 18 training sessions. This measure was estimated only for the experimental group. (a) Boxplots represent values of mean n-level achieved during 18 sessions of training. Error bars represent 95% confidence intervals. On average, participants improved their initial performance by 60.3%. Maximum n-back levels achieved by participants varied from 3-back to 7-back. (b) Growth model fitted to mean n-values. We fitted both linear and quadratic models to predict the behavioral score (mean n-back level) monitored across the 18 training sessions. Training session significantly predicted mean n-back level achieved by participants, *χ*^2^(2) = 111.21, *p* < 0.0001. Including a quadratic term in the model based on session significantly improved the model fit, *χ*^2^(1) = 24.12, *p* < 0.0001. Orange lines represent models of behavioral improvement fitted to each participant’s performance. The black line represents the prototype model fitted to the experimental group. Source data are provided as a Source Data file.

**Supplementary Figure 2:**
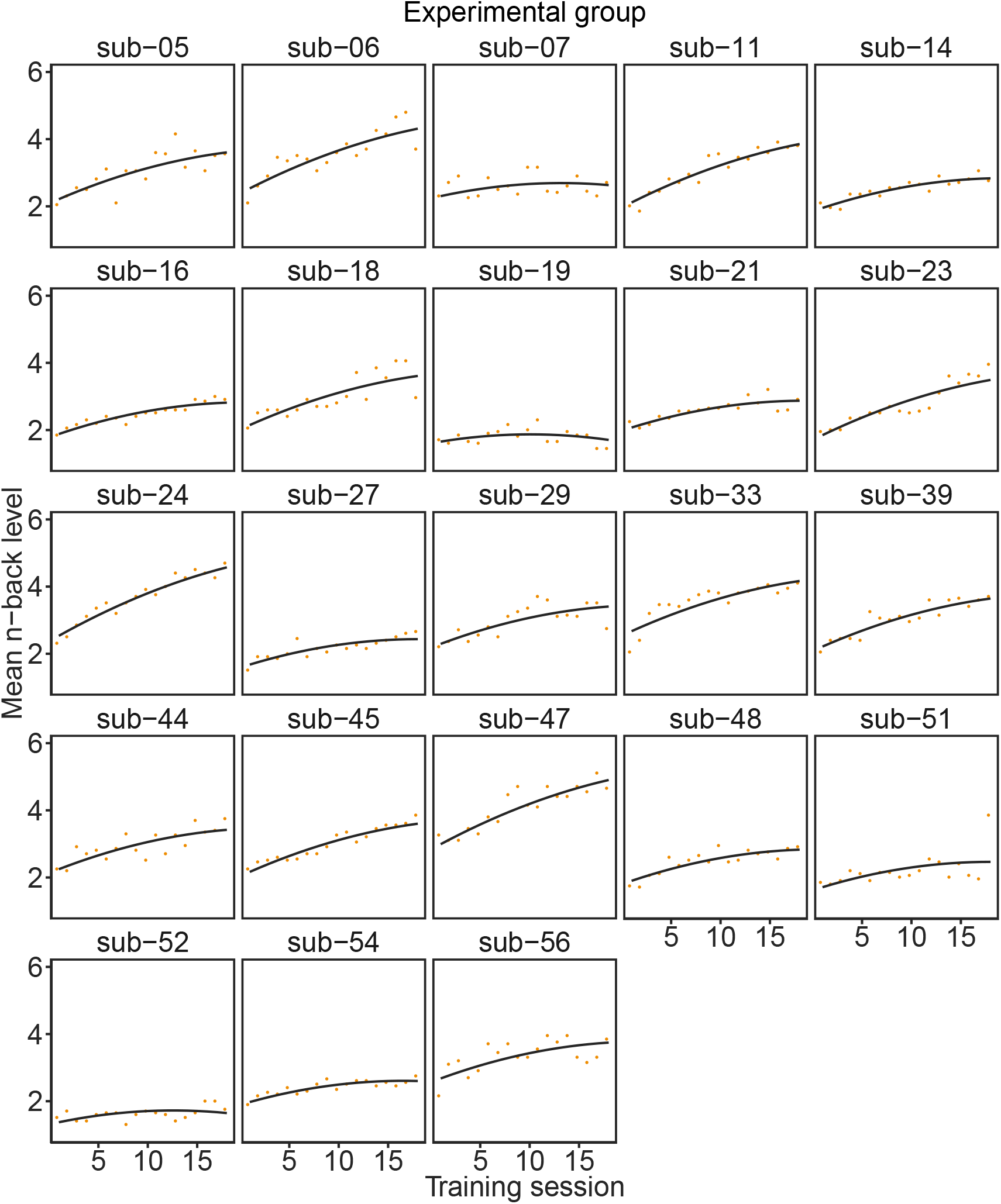
Individual values of mean n-back level achieved in each session of the dual n-back training. The black line represents a quadratic model fitted to individual data. Source data are provided as a Source Data file.

**Supplementary Figure 3:**
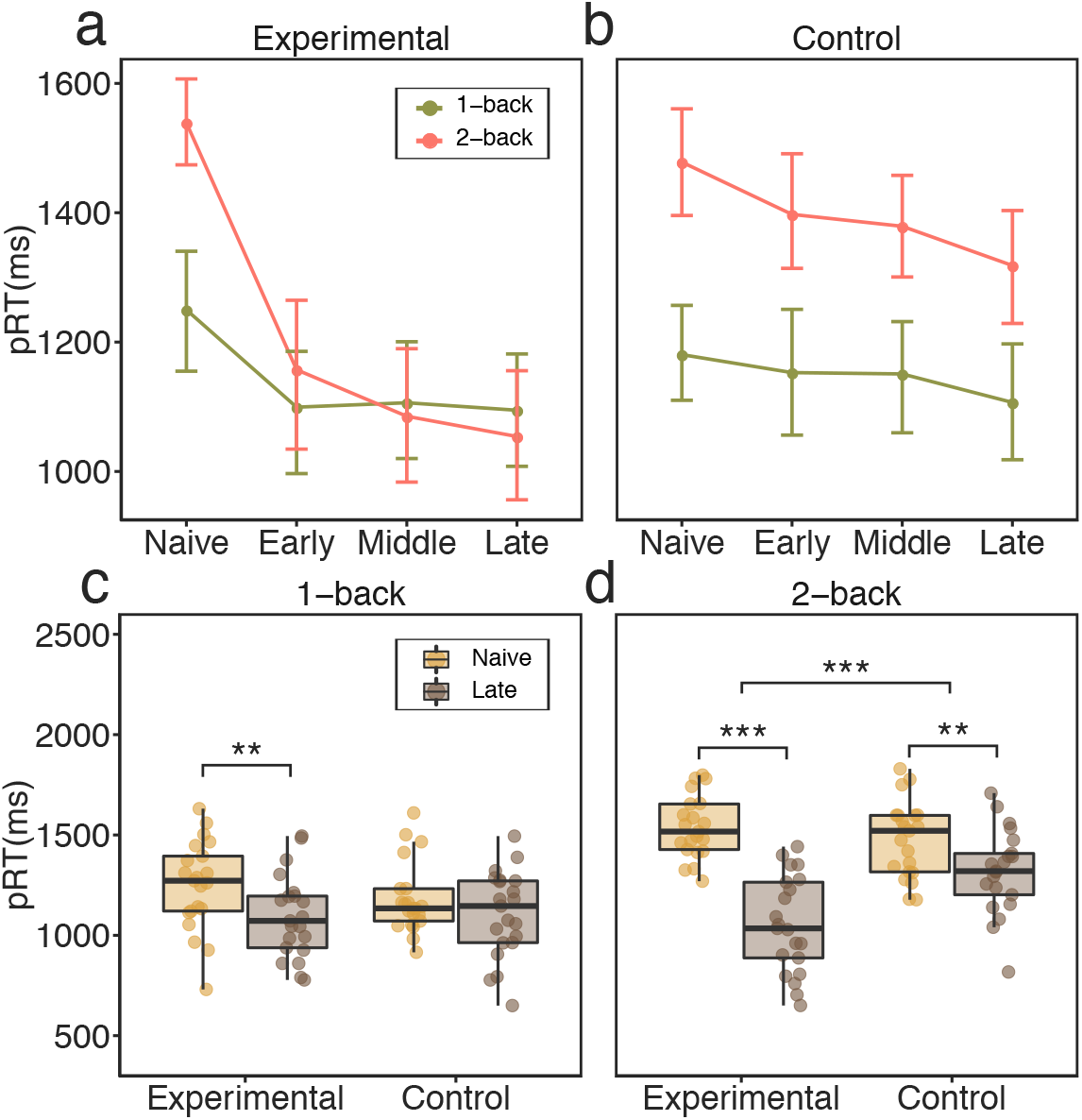
Behavioral performance modulated by training. (**a, b**) Line plots represent mean behavioral performance measured as pRT, calculated for all training phases (Naive, Early, Middle, Late), dual n-back conditions (1-back and 2-back), and groups (experimental, (**a**); control, (**b**). Participants exhibited significantly different pRT, depending on the training stage (Naive, Early, Middle, Late), condition (1-back versus 2-back), and group (experimental versus control), as indicated by a *χ*^2^-test (*χ*^2^(3) = 21.25, *p* < 0.0001). (**c**) In the 1-back condition, the experimental group displayed a 14.2% reduction in pRT (Bonferroni-corrected, *t*(20) = 3.90, *p* = 0.003); no improvement was found in the control group (*t*(20) = 1.77, *p* = 0.08). The change in pRT during the 1-back condition did not differ between the two groups (*t*(39.91) = 1.41, *p* = 0.17). (d) The greatest improvement was observed in the experimental group when comparing ‘Naive’ to ‘Late’ training phases during the 2-back condition (mean 46 % pRT improvement; Bonferroni-corrected, *t*(20) = 10.16, *p* < 0.0001). For comparison, the control group exhibited only a 12.2 % decrease of pRT during the 2-back condition (Bonferroni-corrected, *t*(20) = 3.95, *p* = 0.003). The decrease in pRT was significantly larger for the experimental group than for the control group (Bonferroni-corrected, *t*(38.95) = 5.19, *p* < 0.0001). After training, the experimental group exhibited no difference in performance between the 1-back condition and the 2-back condition after training (t(20) = 1.52, *p* = 0.14), while in control group, the difference in performance between conditions remained substantial (Bonferroni-corrected, *t*(20) = −5.71, *p* < 0.001). Error bars represent 95% confidence intervals. *** *p* < 0.001 Bonferroni-corrected; ** *p* <0.01 Bonferroni-corrected. Source data are provided as a Source Data file.

**Supplementary Figure 4:**
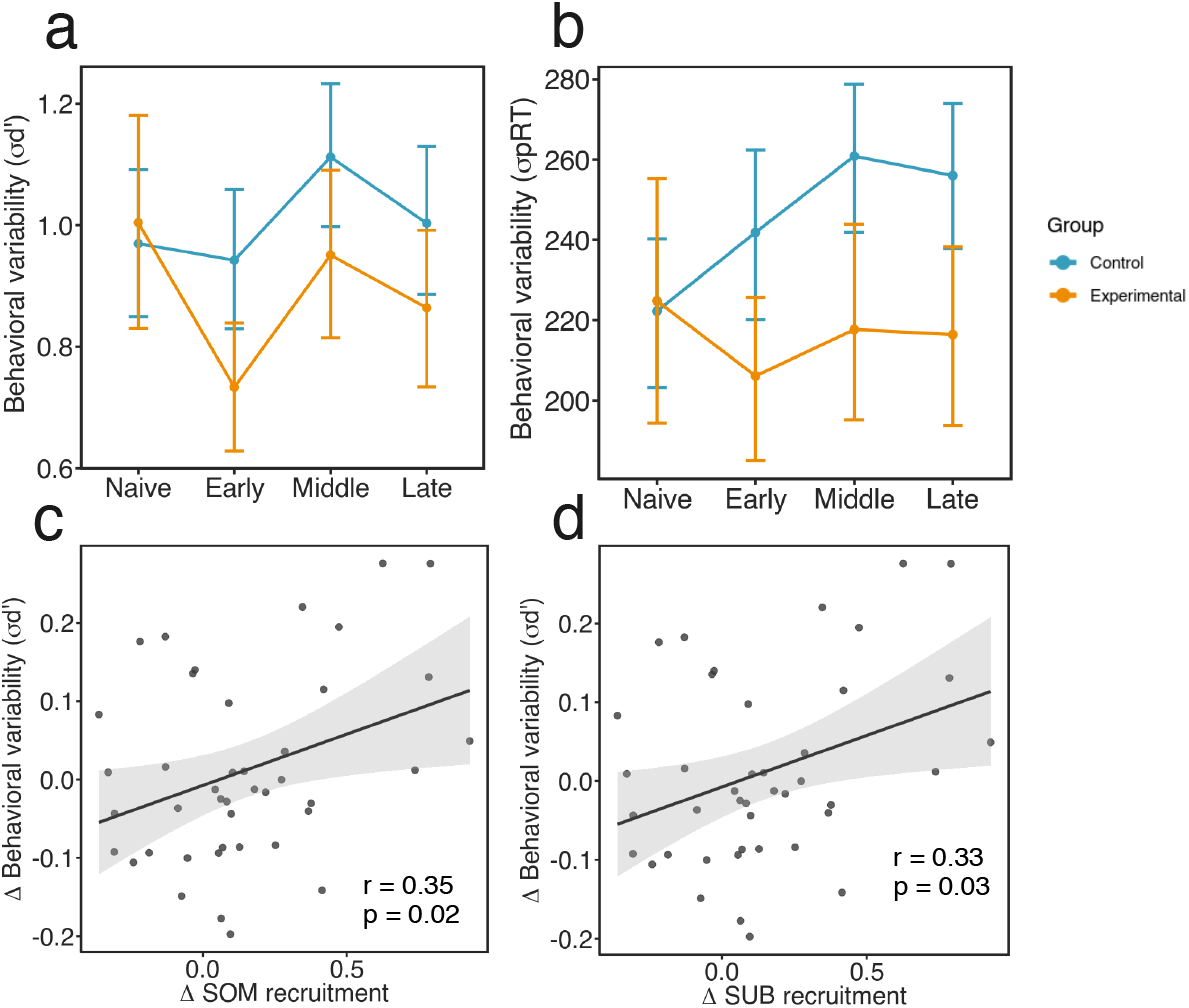
Block-to-block variability in behavioral performance modulated by training. (a) Standard deviation of *d*′ (*σ*_*d*′_) estimated across task blocks, for which we found a significant main effect of session (*χ*^2^(3) = 9.61, *p* = 0.02). Specifically, the standard deviation of *dĩ* decreased from ‘Naive’ to ‘Early’ sessions for all participants (*β* = −0.14, *t*(39) = −2.46, *p* = 0.02). Both group and session × group interaction effects were not significant (*p* > 0.05). Error bars represent the 95% confidence intervals. (b) Standard deviation of penalized reaction time (*σ_pRT_*), for which we found a significant group effect effect(*χ*^2^(1) = 7.39, *p* = 0.006). In general, participants from the experimental group had lower pRT variability (*β*= −29.00, *t*(40) = −2.80, *p* = 0.008) than participants from the control group. Both the effect of session and the session × group interaction were not significant (p > 0.05). (c, d) correlation between the across-session change in behavioral variability measured as standard deviation of *d*′ and the across-session change in (c) somatomotor and (d) subcortical systems recruitment (*p* < 0.05, uncorrected). Source data are provided as a Source Data file.

**Supplementary Figure 5:**
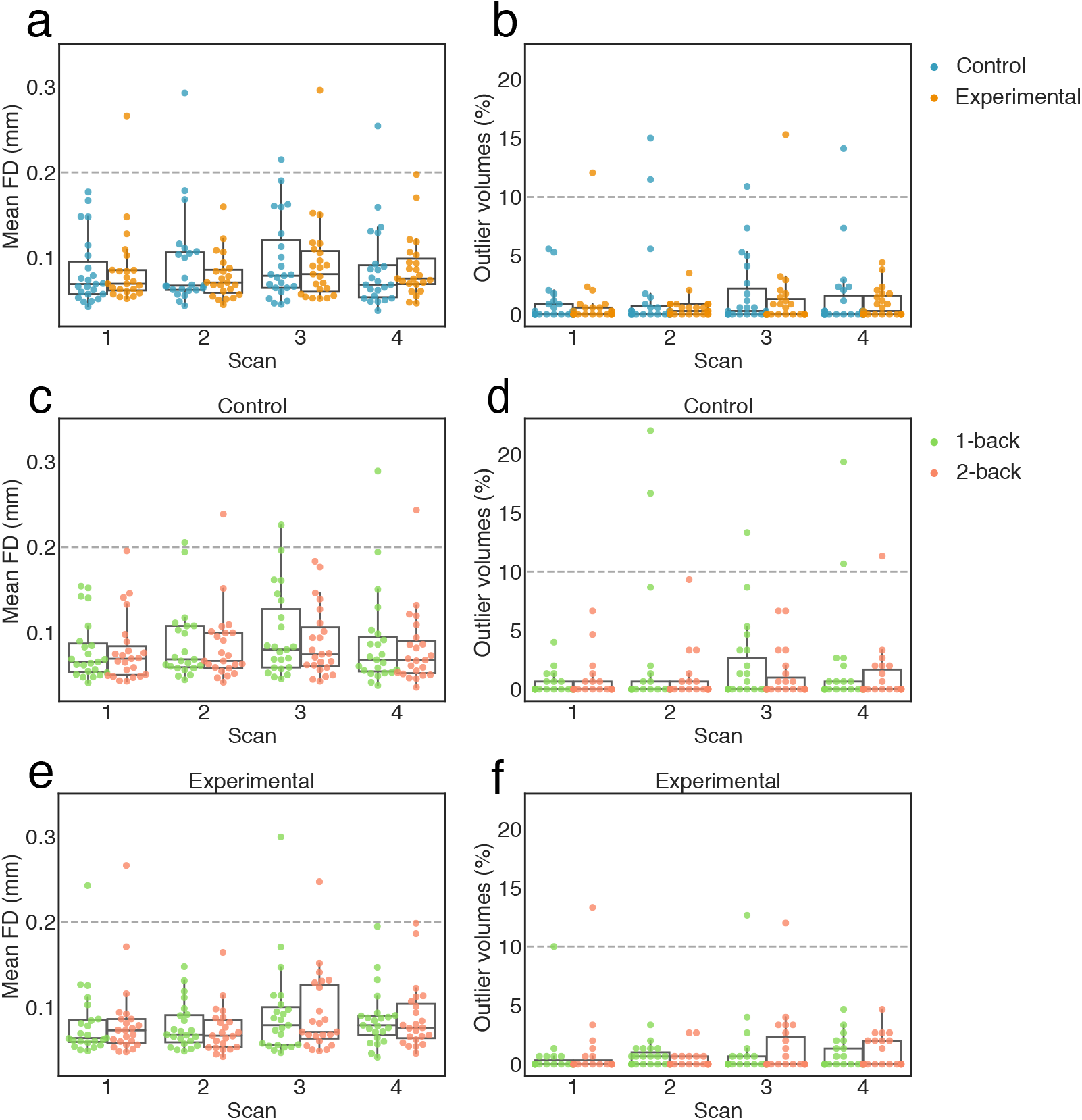
Head motion during the dual n-back task. In addition to including 24 motion parameters in the denoising procedure, we also excluded high motion subjects from subsequent analyses. We defined a high motion subject as one with mean frame displacement (FD) larger than 0.2 mm and more than 10% of outlier volumes in any scanning session. This criterion was applied when considering the total time courses, as well as when considering time courses of the 1-back and 2-back conditions, separately. As a result we excluded four participants (2 from the control group, and 2 from the experimental group). One subject displayed excessive motion during three scanning sessions, while another displayed excessive motion during two scanning sessions, and two subjects displayed excessive motion in only one scanning session. After excluding high motion subjects, we compared the mean FD and mean percent of outlier scans between sessions, groups, and conditions (see S1 for further details). We did not find significant differences between any of these variables between sessions (all *p* < 0.05), groups (all < 0.05), and most of the condition comparisons. The only difference that passed an uncorrected threshold of significance (*p* < 0.05) was found between the 1-back and 2-back conditions of the control group during the third scanning session. Source data are provided as a Source Data file.

**Supplementary Figure 6:**
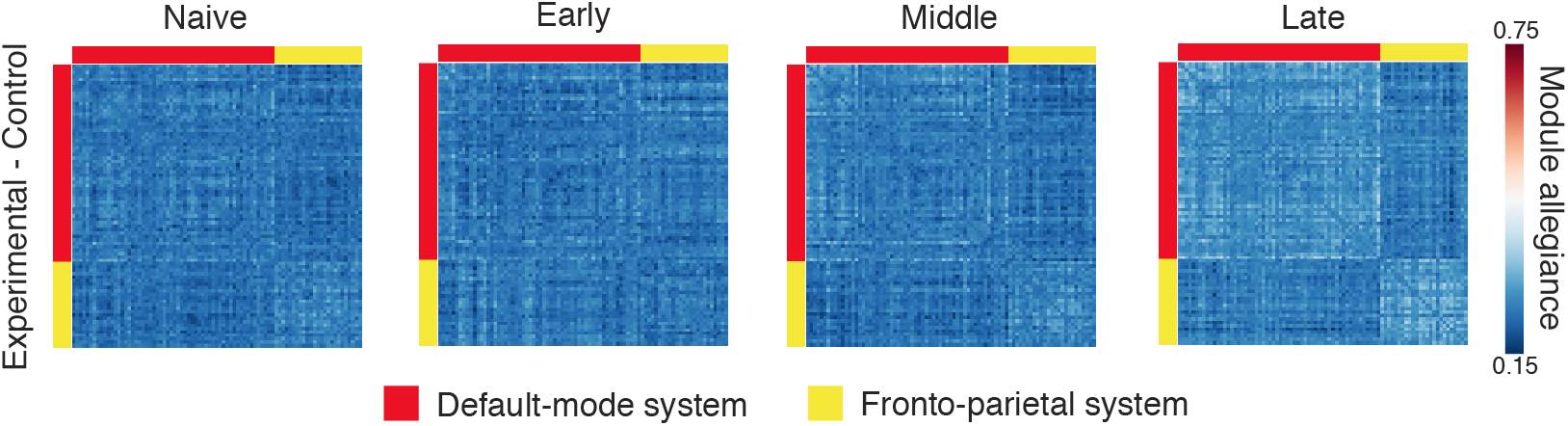
Between-group differences in module allegiance matrices for the default mode and frontoparietal systems. Each *ij*-th element of the matrix represents a difference between groups (experimental minus control) in the probability that node *i* and node *j* are assigned to the same module within a single layer of the multilayer network. Systems are defined using the Power et al.^1^ parcellation. Source data are provided as a Source Data file.

**Supplementary Figure 7:**
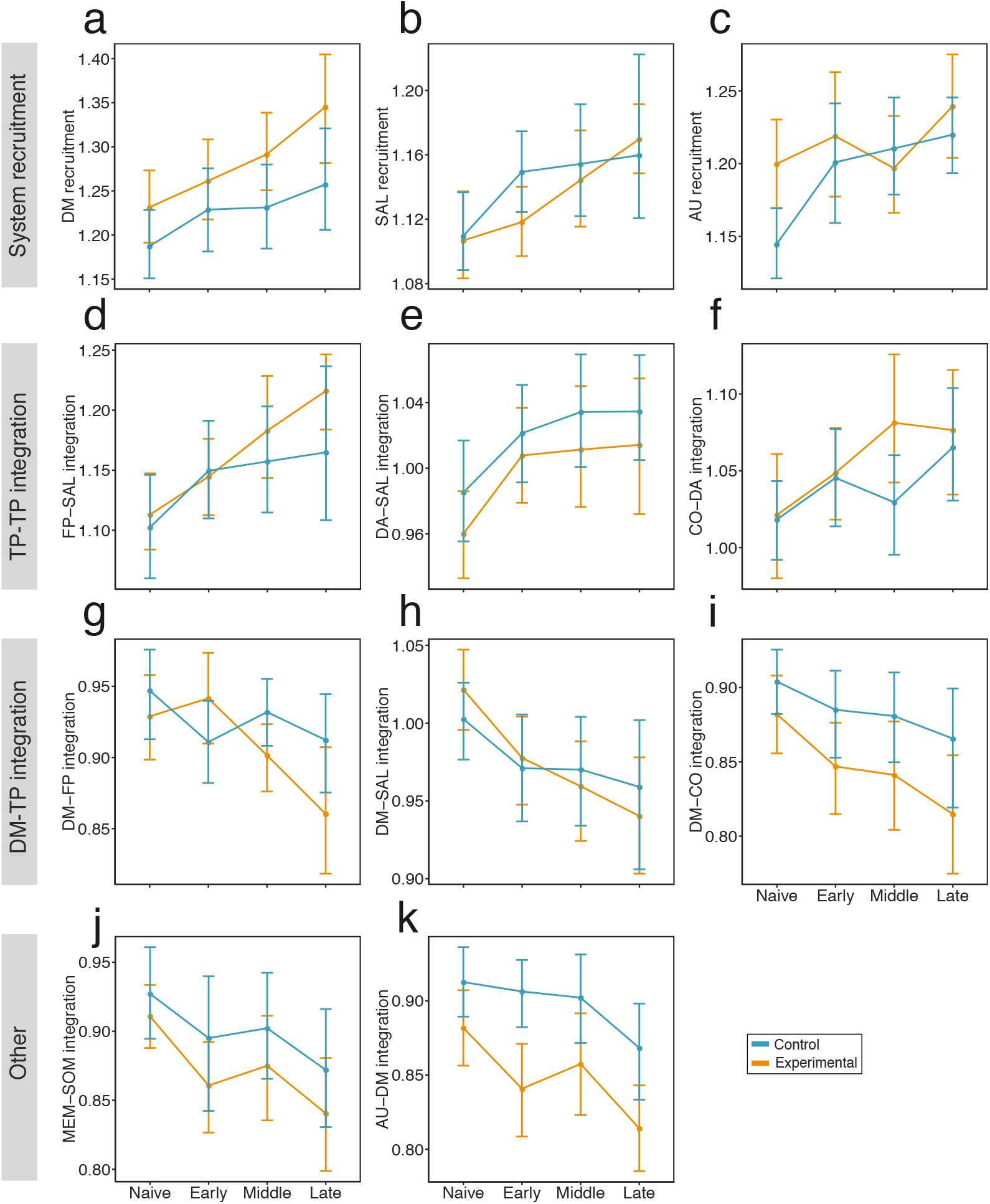
Session-to-session changes in recruitment and integration of large-scale systems. We observed three main categories of large-scale system reorganization: (**a-c**) an increase in system recruitment, (**d-f**) an increase in integration between task-positive systems (TP), (**g-i**) a decrease in integration between the default mode (DM) system and task-positive systems, and (**j-k**) other. Error bars represent 95% confidence intervals. Remaining abbreviations: salience (SAL), auditory (AU), fronto-parietal (FP), dorsal attention (DA), cingulo-opercular (CO), memory (MEM), and somatomotor (SOM). Source data are provided as a Source Data file.

**Supplementary Figure 8:**
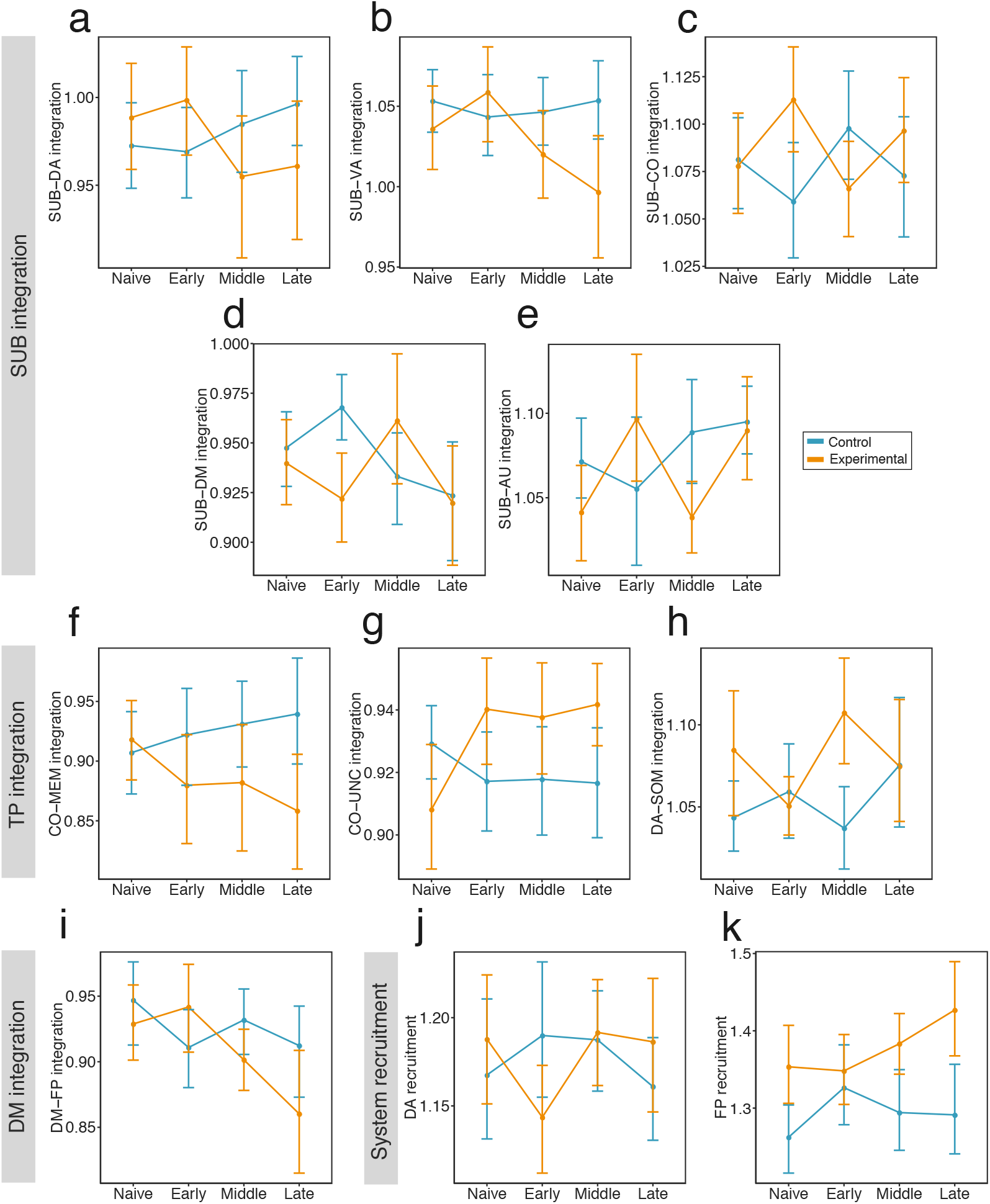
Session × group interaction effects for across-session changes in recruitment and integration of large-scale systems. We observed group differences in the changes in (**a-e**) integration of subcortical (SUB) system with other systems, (**f-h**) integration of task-positive (TP) systems with other systems, (**i**) integration of the default mode (DM) with the fronto-parietal (FP) system, (**j-k**) changes in dorsal attention (DA) and FP recruitment. Error bars represent 95% confidence intervals. Remaining abbreviations: salience (SAL), auditory (AU), memory (MEM), and somatomotor (SOM). Source data are provided as a Source Data file.

**Supplementary Figure 9:**
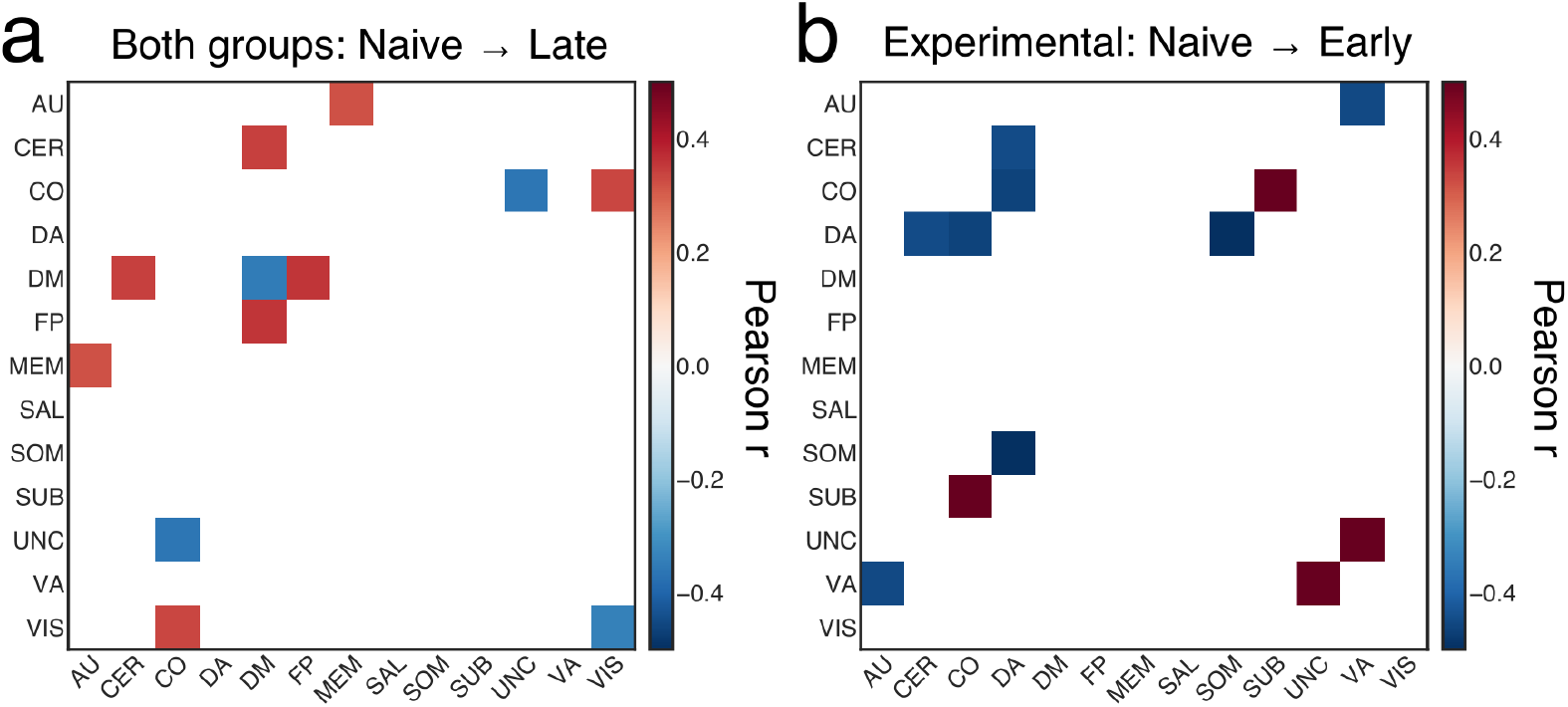
Relationship between the change in network dynamics and the change in behavior. Colored tiles represent all correlations (p < 0.05, uncorrected; \**p* < 0.05 FDR-corrected). (**a**) Pearson correlation coefficient (*r*) between the across-session changes in recruitment (or integration) and the across-session changes in penalized reaction time (Δ pRT) observed for both experimental and control group. (**b**) Relationship between the changes in recruitment (or integration) and the changes in pRT during early phase of training of the experimental group. Abbreviations: auditory (AU), cerebellum (CER), cingulo-opercular (CO), default mode (DM), dorsal attention (DA), fronto-parietal (FP), memory (MEM), salience (SAL), somatomotor (SOM), subcortical (SUB), uncertain (UNC), ventral attention (VA), and visual (VIS). Source data are provided as a Source Data file.

**Supplementary Figure 10:**
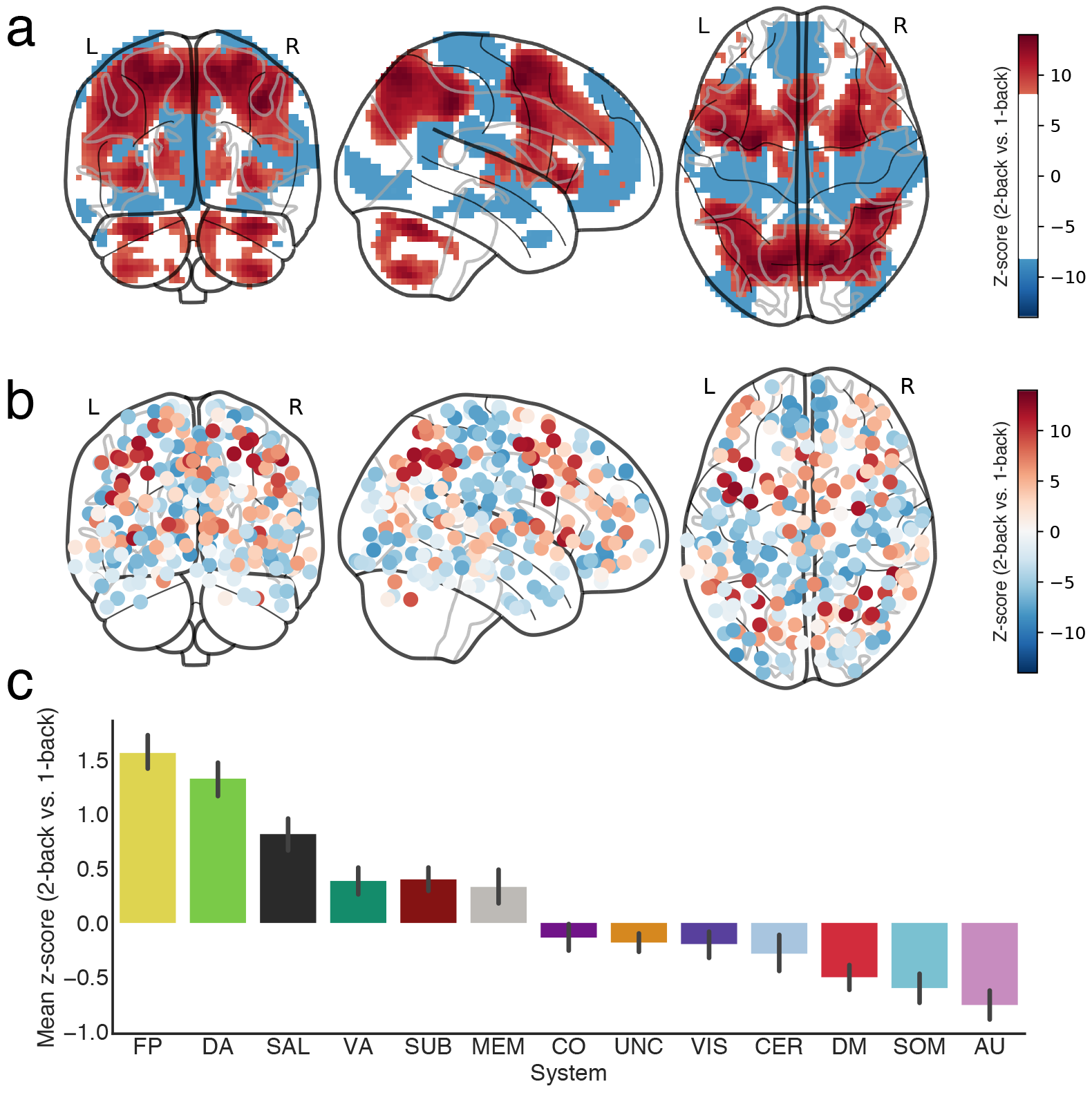
Brain activity for 2-back vs. 1-back contrast (two-sided) estimated with a standard GLM for all subjects and sessions. (**a**) Glass brain visualization of activity thresholded at a z-score level ±8. (b) Brain activity plotted on 264 ROIs from the Power et al.^1^ parcellation. (**c**) Barplot representing the z-score values averaged over ROIs belonging to predefined large-scale systems. The most active ROIs belonged to the fronto-parietal (FP), dorsal attention (DA), and salience systems (SAL). The most deactivated ROIs belonged to the auditory (AU), somatomotor (SOM), and default mode (DM) systems. Remaining abbreviations: cerebellum (CER), cingulo-opercular (CO), memory (MEM), uncertain (UNC), somatomotor (SOM), subcortical (SUB), ventral attention (VA), and visual (VIS). Source data are provided as a Source Data file. Source data are provided as a Source Data file.

**Supplementary Figure 11:**
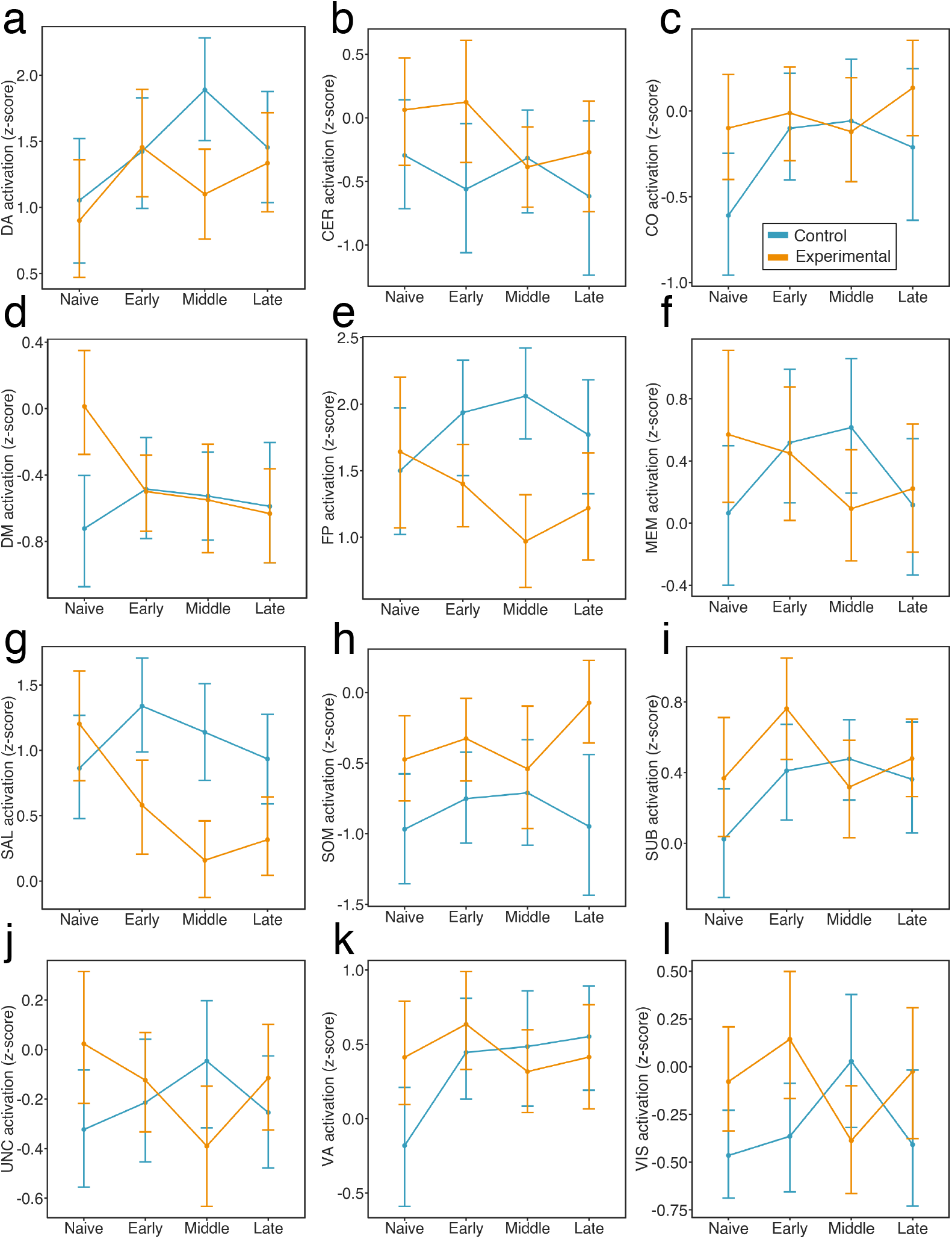
Cross-sessions changes in brain activity for 2-back vs. 1-back contrast (two-sided) estimated with a standard GLM. Groups differed significantly by session for the salience (SAL) and visual (VIS) systems (*p* < 0.05, FDR-corrected). Specifically, compared to the control group, participants from the experimental group displayed significantly greater decreases in the activation of the salience system from the ‘Naive’ to ‘Early’ sessions (*β* = −1.10, *t*(120) = −3.44, *p* = 0.0008), from the ‘Naive’ to ‘Middle’ sessions (*β* = −1.32, *t*(120) = −4.12, *p* = 0.0001), and from the ‘Naive’ to ‘Late’ sessions (*β* = −0.96, *t*(120) = −2.99, *p* = 0.003). The experimental group also displayed a larger decrease in the activation of the visual system from the ‘Naive’ to ‘Middle’ sessions: *β* = −0.80, *t*(120) = −2.69, *p* = 0.008). Source data are provided as a Source Data file.

**Supplementary Figure 12:**
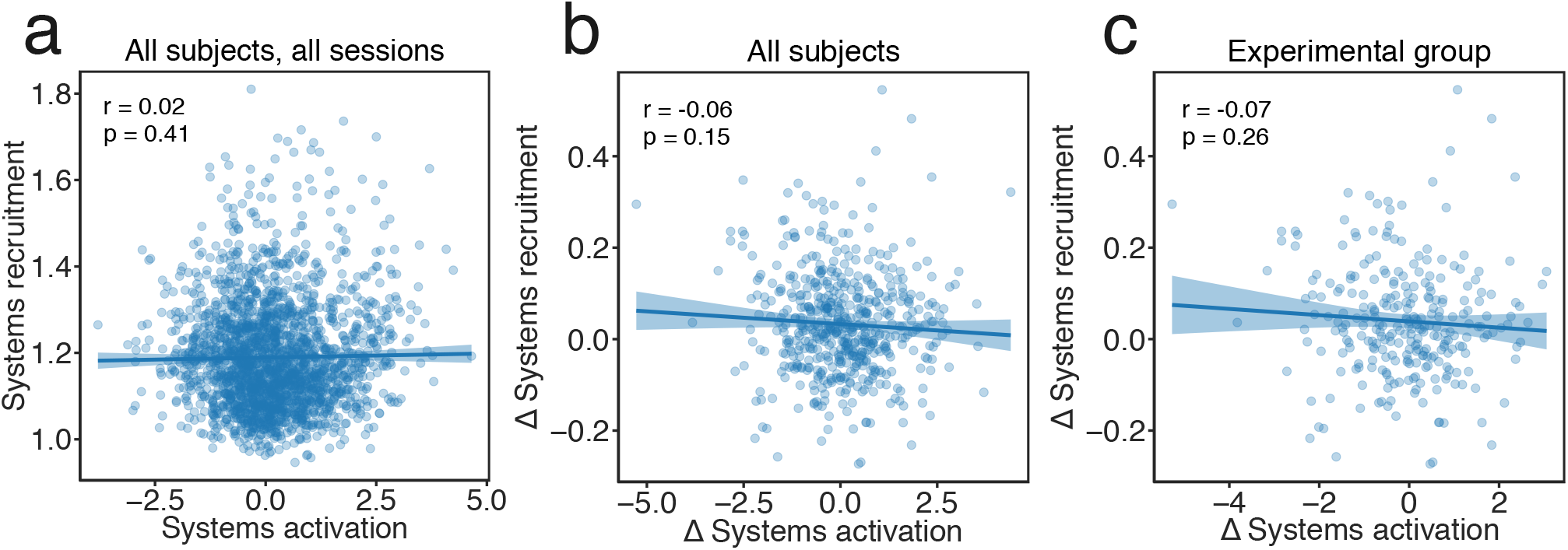
Relationship between systems recruitment and systems activation estimates. (**a**) There was no significant relationship between systems activation (z-score; 2-back minus 1 back) and systems recruitment values when considering all systems, all sessions, and all subjects. We further tested whether changes (*δ*) in systems activity from ‘Naive’ to ‘Late’ sessions were correlated with changes in systems recruitment. We did not find any significant correlation between these two variables, either when considering (**b**) all subjects, or (**c**) when considering only the experimental group. Source data are provided as a Source Data file.

**Supplementary Figure 13:**
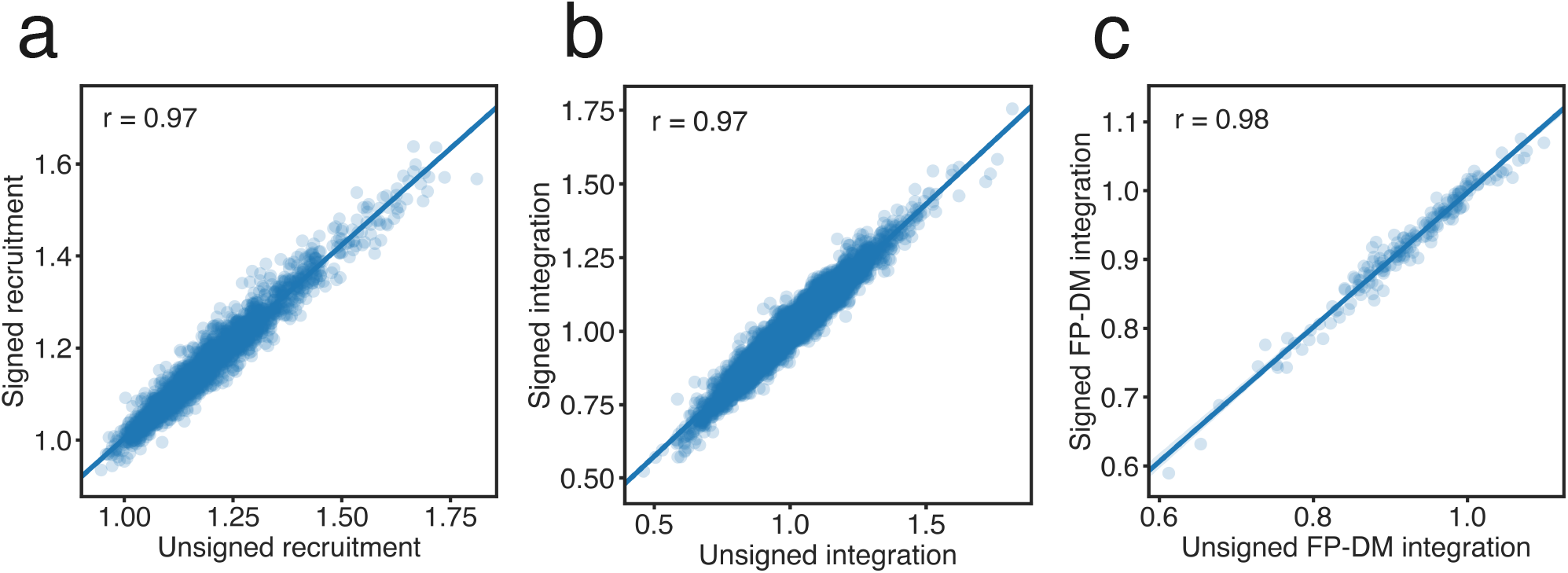
Relationship between recruitment and integration values calculated based on unsigned and signed functional connectivity matrices. Unsigned and signed recruitment (**a**) and integration (**b**) coefficients estimated for all large-scale systems were highly correlated. (**c**) Values of integration between fronto-parietal (FP) and default mode (DM) systems were also highly correlated. Source data are provided as a Source Data file.

**Supplementary Figure 14:**
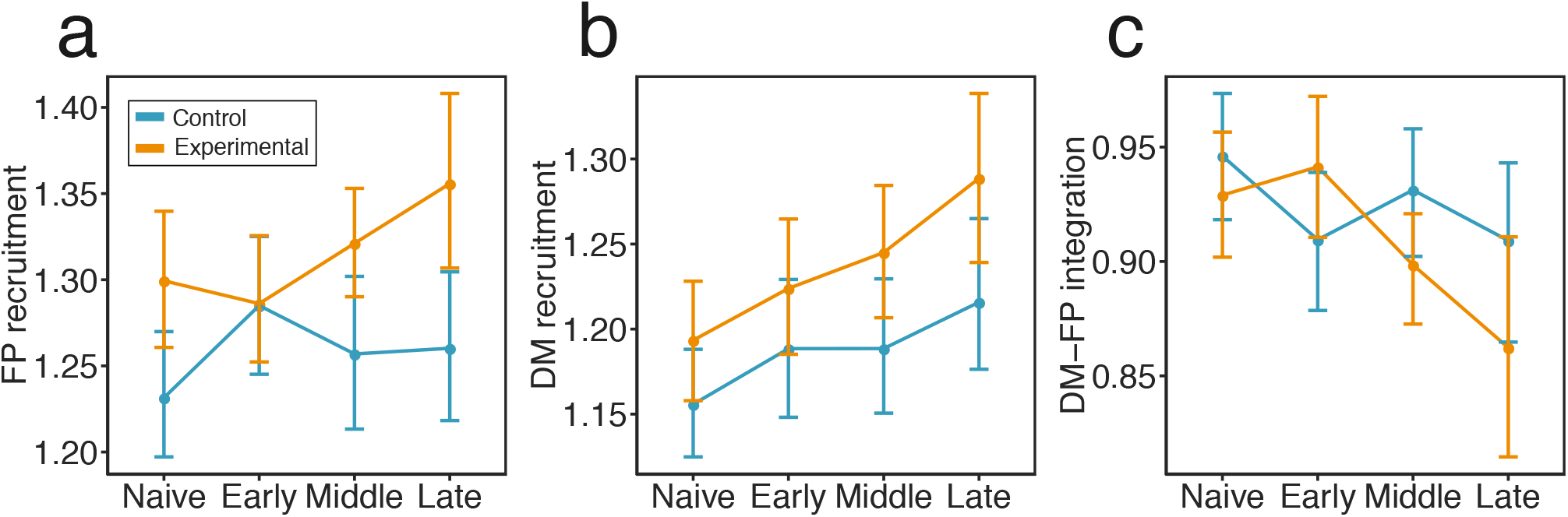
Changes in module allegiance of the fronto-parietal (FP) and default-mode (DM) systems calculated based on signed functional connectivity matrices. We observed a significant session × group interaction effect when considering changes in the recruitment of the fronto-parietal system during training (*χ*^2^(3) = 9.31, *p* = 0.025) (**a**). The largest increase in fronto-parietal recruitment was observed in the experimental group when comparing ‘Early’ to ‘Late’ training phases (*β* = −0.07, *t*(120) = −3.057, *p* = 0.016, Bonferroni-corrected). No significant changes from ‘Naive’ to ‘Late’ training phases were observed in the control group (*β*= −0.05, *t*(120) = −2.35, *β* = 0.12, Bonferroni-corrected). (**b**) Turning to an examination of the default mode, we found a significant main effect of session (*χ*^2^(3) = 23.89, *p* < 0.0001) on system recruitment However, the interaction effect between session and group was not significant (*χ*^2^(3) = 2.00, *p* = 0.57). Planned contrasts revealed that the default mode recruitment increased steadily in both groups and we observed the largest increase between ‘Naive’ and ‘Late’ sessions (*β*= 0.08, *t*(123) = 5.02, *p* < 0.0001). (**c**) We found a significant session × group interaction effect on the integration between the fronto-parietal and default mode systems (*χ*^2^(3) = 13.30, *p* = 0.004). The integration between these two systems decreased from ‘Early’ to ‘Late’ sessions only in the experimental group (*β* = 0.08, *t*(120) = 4.86, *p* = 0.0035, Bonferroni-corrected). However, groups differed from ‘Naive’ to ‘Early’ (*β* = 0.05, *t*(120) = 2.13, *p* = 0.03) and from ‘Early’ to ‘Middle’ sessions (*β* = −0.06, *t*(120) = −2.81, *p* = 0.02. Source data are provided as a Source Data file.

**Supplementary Figure 15:**
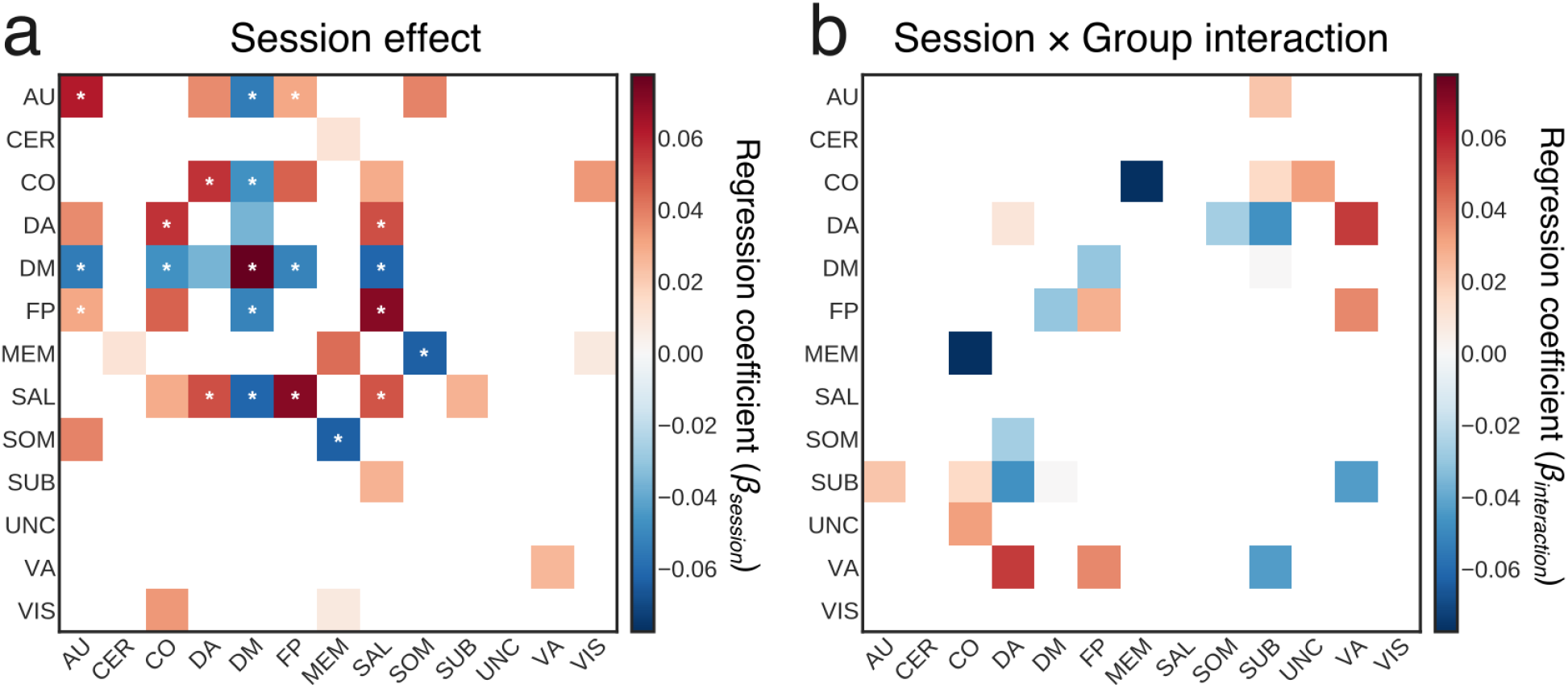
Changes of the recruitment and integration of large-scale systems calculated based on signed functional connectivity matrices. Colored tiles represent all significant effects (*p* < 0.05, uncorrected; \**p* < 0.05 FDR-corrected). (**a**) Here we display the significant main effects of session. Tile color codes a linear regression coefficient (*β*), for all main session effects (from ‘Naive’ to ‘Late’). (**b**) Here we display the significant session group interaction effects. Tile color codes a linear regression coefficient between groups and sessions (from ‘Naive’ to ‘Late’). Abbreviations: auditory (AU), cerebellum (CER), cingulo-opercular (CO), default mode (DM), dorsal attention (DA), fronto-parietal (FP), memory (MEM), salience (SAL), somatomotor (SOM), subcortical (SUB), uncertain (UNC), ventral attention (VA), and visual (VIS). Source data are provided as a Source Data file.

**Supplementary Figure 16:**
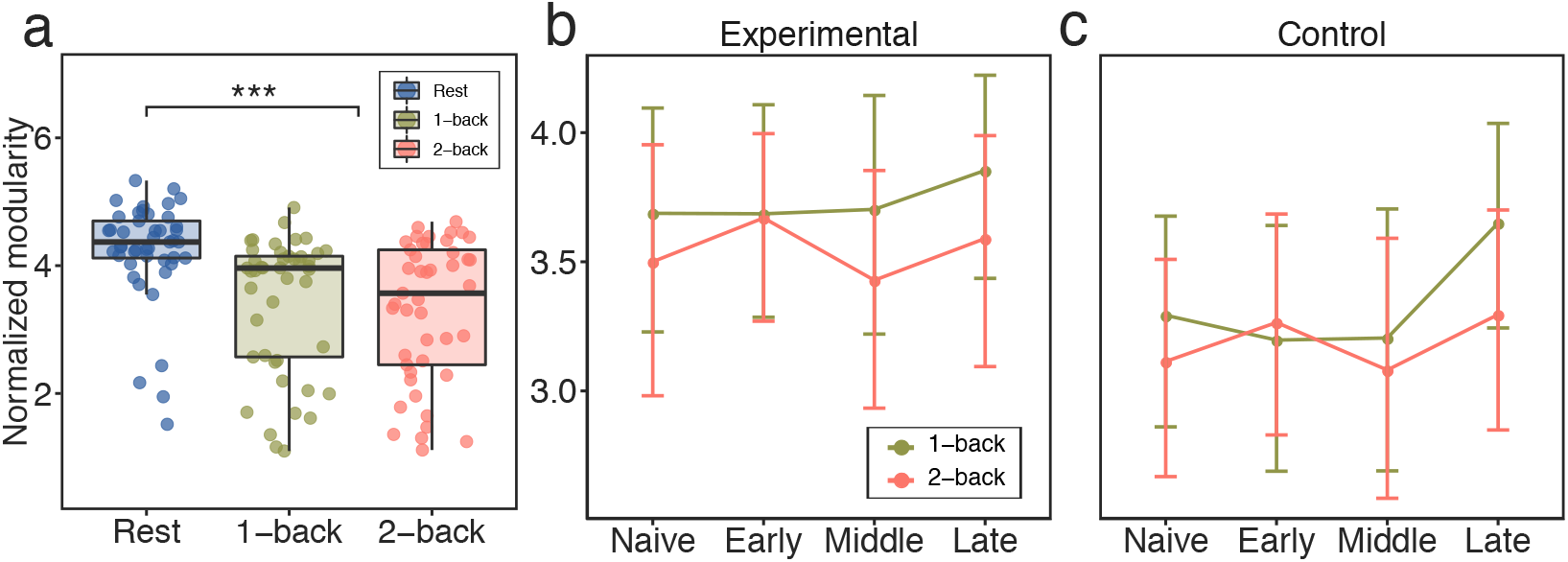
Whole-brain modularity obtained for the Schaefer parcellation. (a) Modularity differences between the resting state and the dual n-back task, as well as between the 1-back task condition and the 2-back task condition. (**b, c**) Line plots representing mean values of modularity for each scanning session (Naive, Late, Middle, Late) and condition, separately for the experimental group (b) and for the control group (c). Source data are provided as a Source Data file.

**Supplementary Figure 17:**
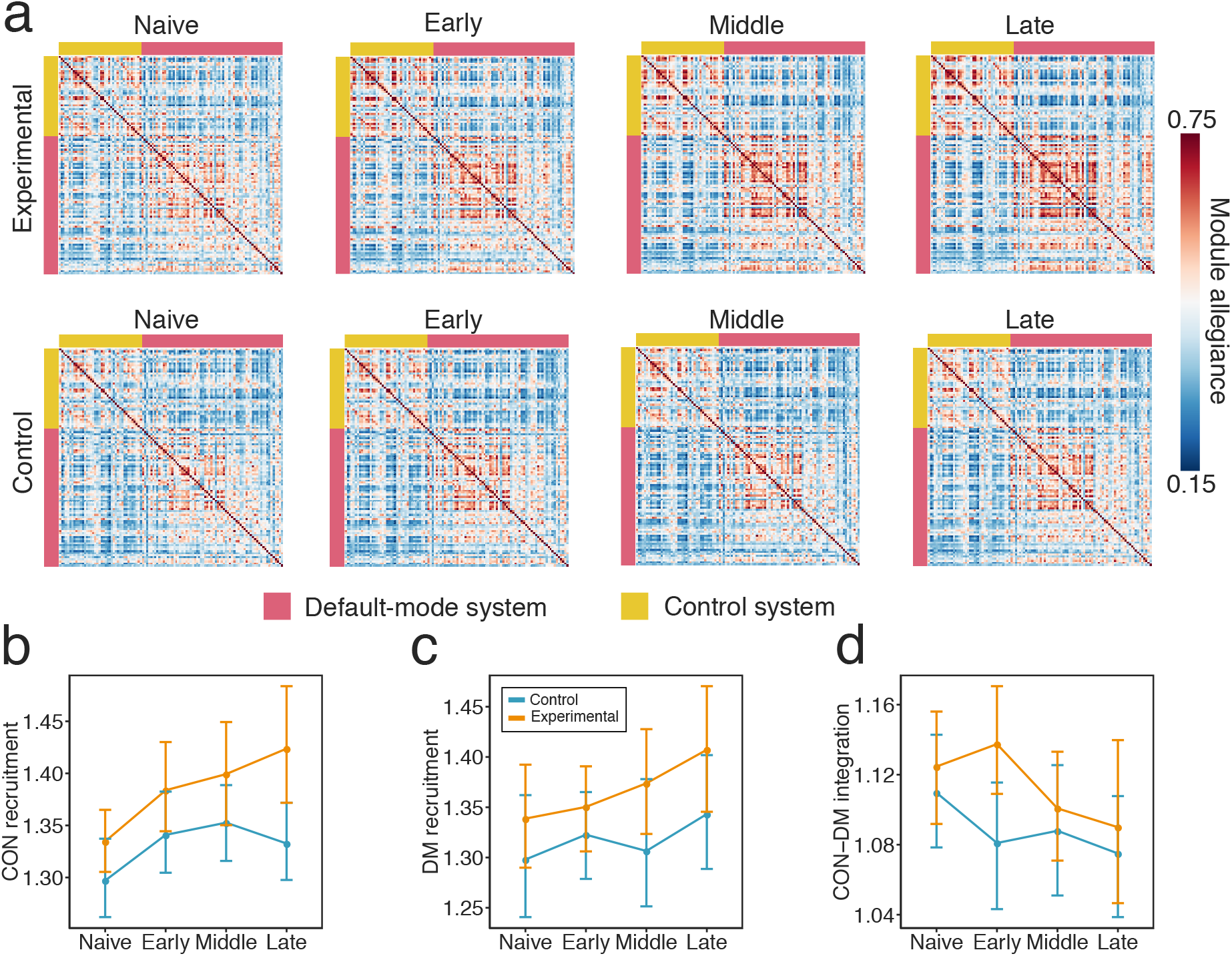
Training-related changes in module allegiance for the subgraph of the network composed of the default mode and fronto-parietal control (CON) systems calculated using the Schaefer parcellation. (**a**) Module allegiance matrices of the default mode system and the fronto-parietal control system (CON). Each *ij*-th element of the module allegiance matrix represents the probability that node *i* and node *j* are assigned to the same community within a single layer of the multilayer network representing task conditions pooled across all scanning sessions. (**b**) Mean CON recruitment across sessions. (**c**) Mean default mode system recruitment across sessions. (**d**) Mean integration between the default mode and CON systems across sessions. Only CON recruitment exhibited a significant main effect of session (*p* < 0.002). Source data are provided as a Source Data file.

**Supplementary Figure 18:**
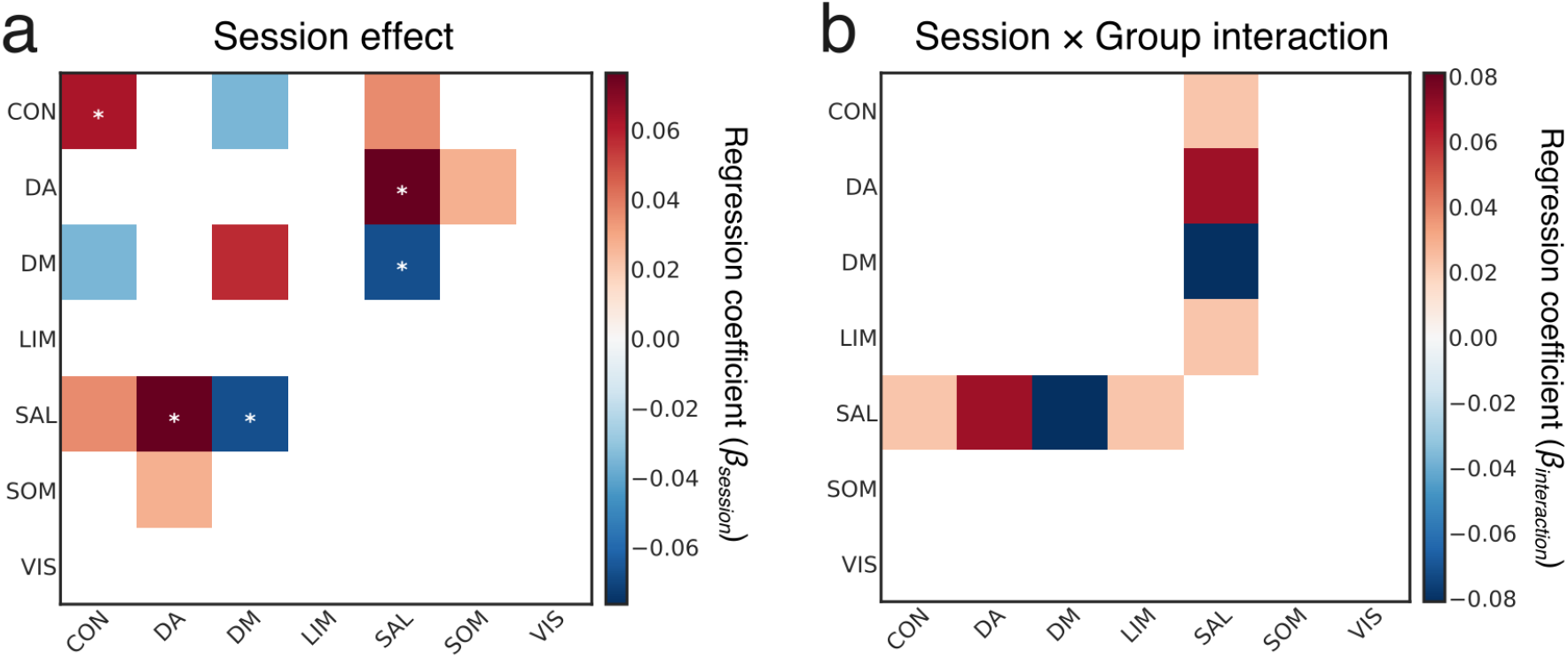
Changes of the recruitment and integration of large-scale systems calculated for Schaefer parcellation^2^. Colored tiles represent all significant effects (*p* < 0.05, uncorrected; \**p* < 0.05 FDR-corrected). (**a**) Here we display the significant main effects of session. Tile color codes a linear regression coefficient (*β*), for all main session effects (from ‘Naive’ to ‘Late’). (**b**) Here we display the significant session × group interaction effects. Tile color codes a linear regression coefficient between groups and sessions (from ‘Naive’ to ‘Late’). Abbreviations: control (CON), dorsal attention (DA), default mode (DM), limbic (LIM), salience (SAL), somatomotor (SOM), visual (VIS). Source data are provided as a Source Data file.

**Supplementary Figure 19:**
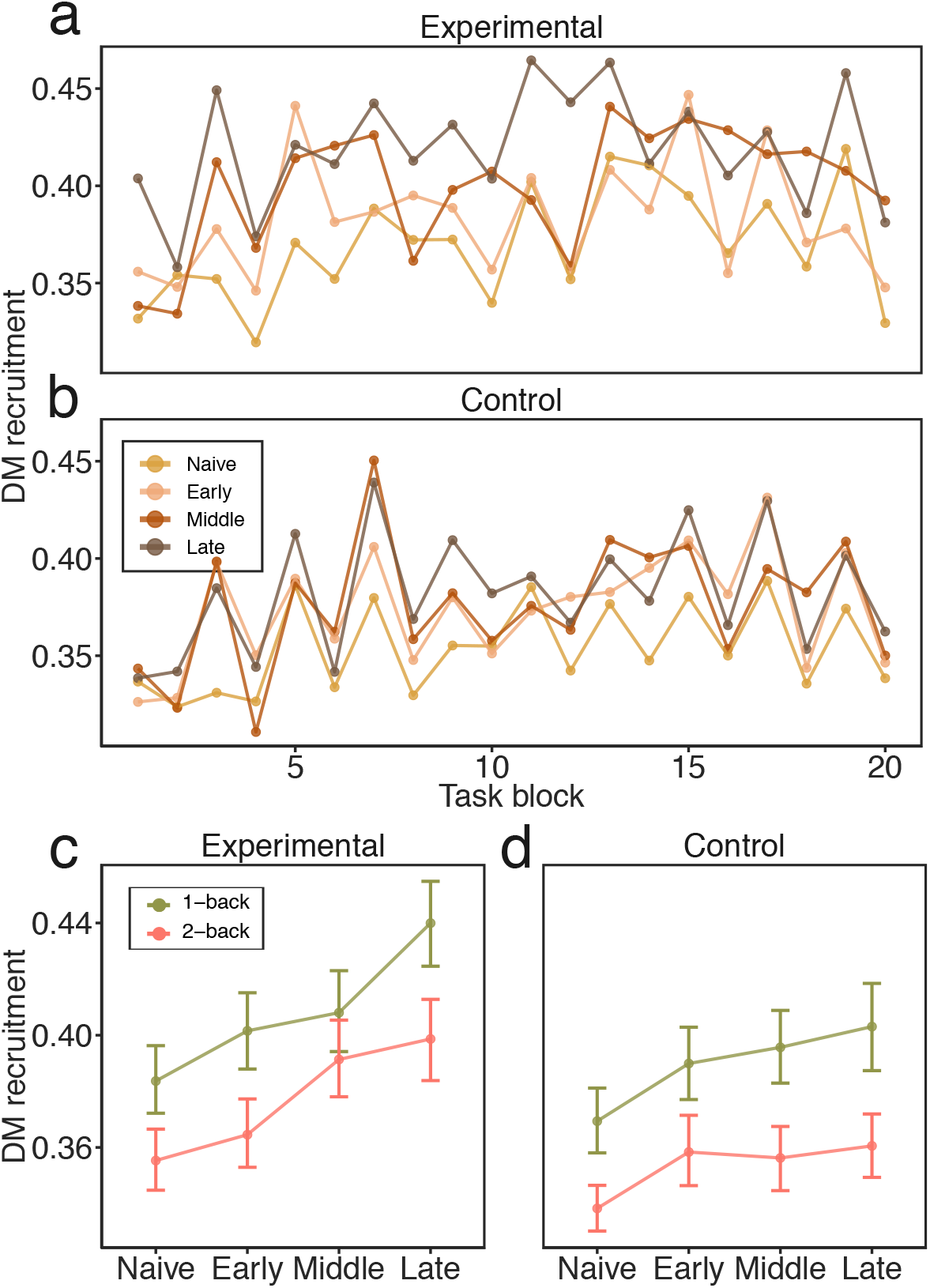
Fluctuations in the recruitment of the default mode system across task blocks. We examined changes in the default mode recruitment by calculating allegiance matrices for each task block. We found a significant effect of condition (*χ*^2^(3) = 83.97, *p* < 0.00001), such that the recruitment of the default mode fluctuated between task conditions and was significantly higher in the 1-back condition (M = 0.40) than in the 2-back condition (M = 0.36; *t*(167) = −10.43, *p* < 0.00001). However, the session × condition interaction was not significant (*χ*^2^(3) = 2.82, *p* = 0.40). Collectively, these results suggest that the default mode recruitment is not only modulated by working memory training, but also by the changing demands of the cognitive task. Across-block fluctuations in default mode recruitment in (**a**) the experimental group and (**b**) the control group. Differences between task conditions for (**c**) the experimental group and (**d**) the control group, across training stages. Source data are provided as a Source Data file.

**Supplementary Figure 20:**
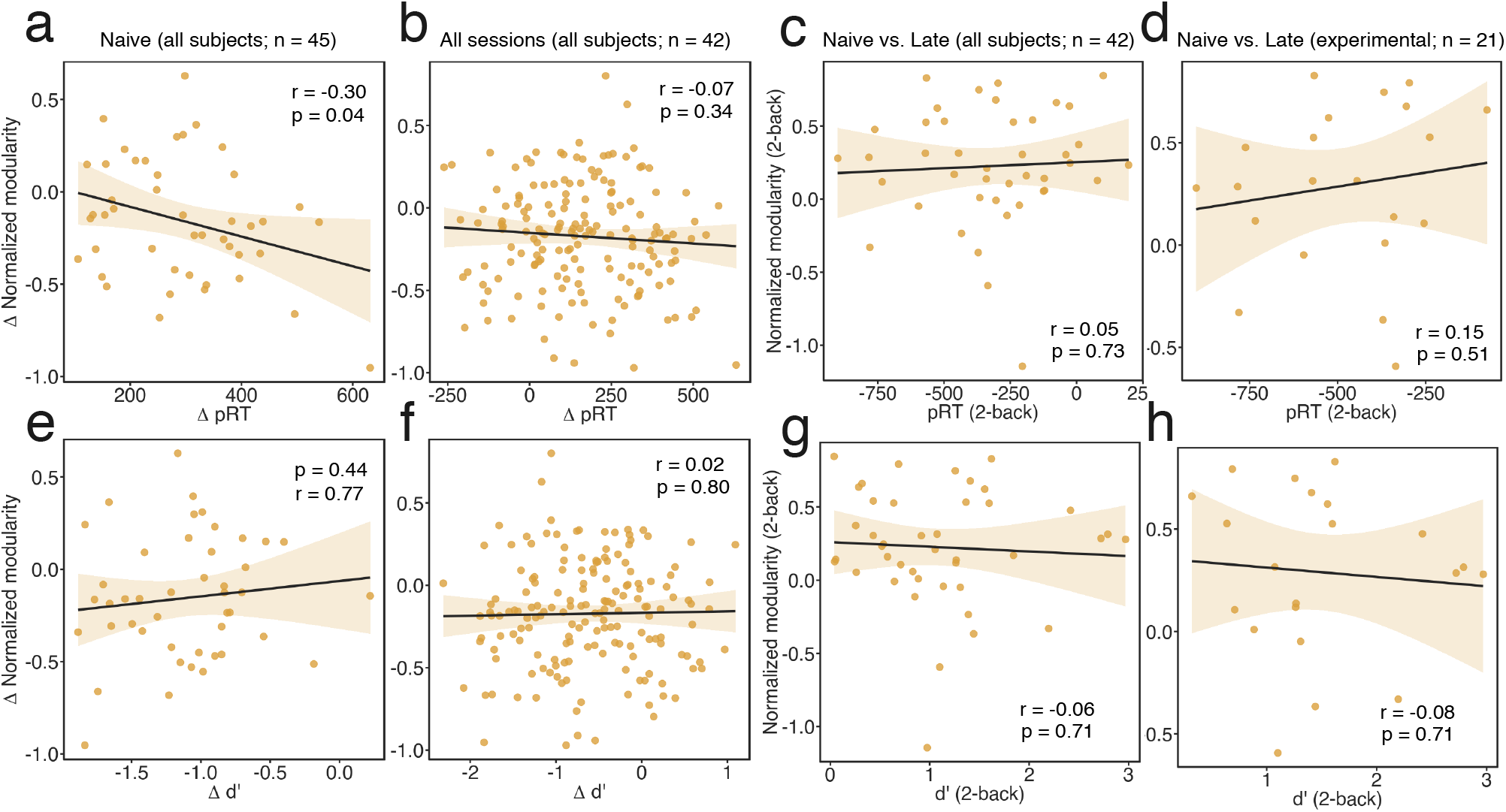
Relationship between modularity and behavioral performance. (a) We observed a weak negative correlation between the change (Δ) of modularity (2-back - 1-back) and the change in penalized reaction time (Δ pRT) during ‘Naive’ session. (b) Change of modularity was not related to the changes in pRT when considered all scanning sessions. (e, f) We did not observe any relationship between the change in d’ and the change in modularity for ‘Naive’ and for all scanning sessions. (c, d, g, h) The change of modularity during 2-back was not correlated to the changes in pRT or *d*′ from ‘Naive’ to ‘Late’ session.

**Supplementary Figure 21:**
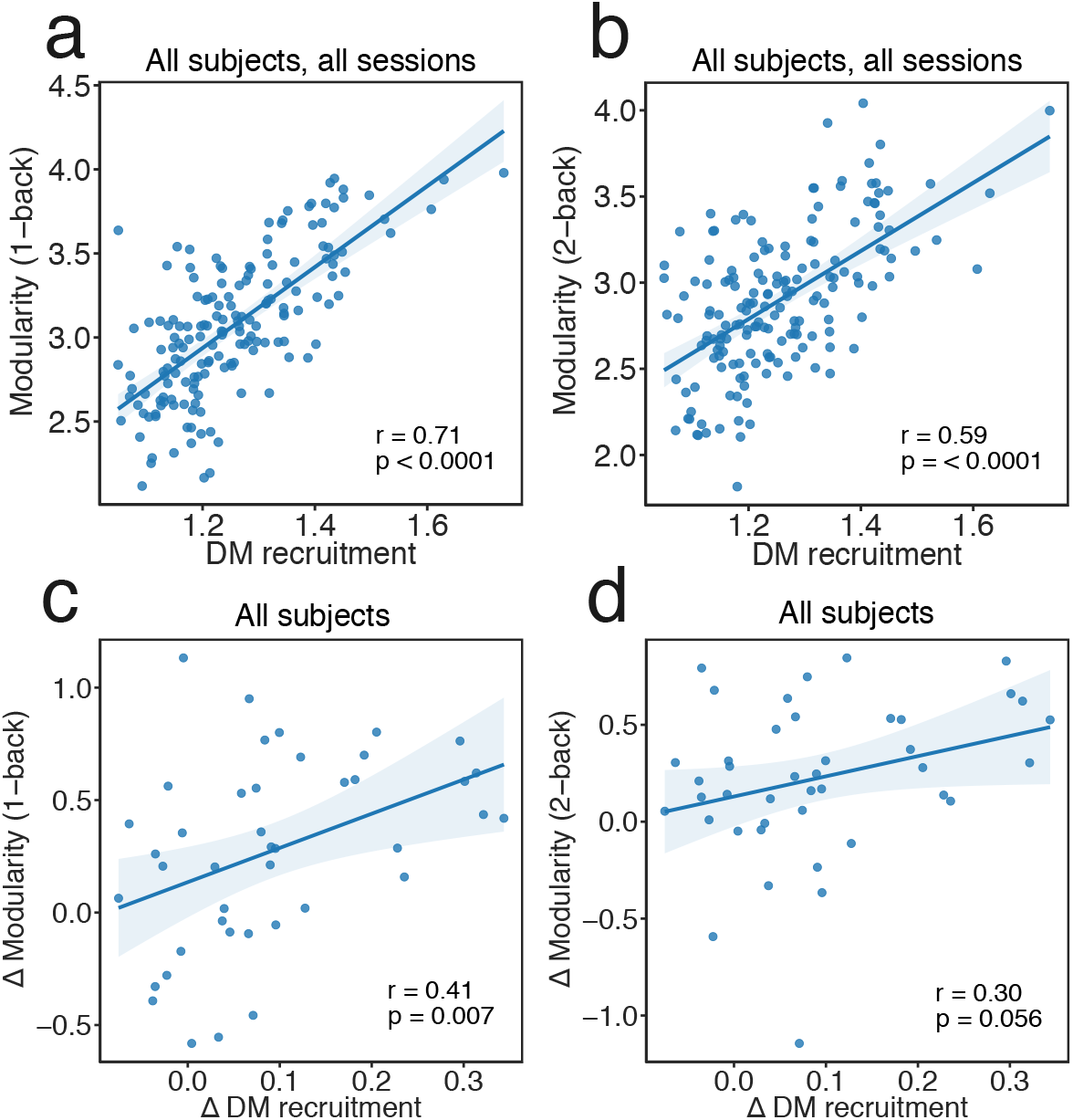
Relationship between default mode recruitment and static modularity. Correlation between DM recruitment and modularity during (**a**) 1-back and (b) 2-back conditions calculated for all subjects and all sessions. Correlation between the change (Δ) of DM recruitment and change of modularity during (c) 1-back condition and (**d**) 2-back condition. Source data are provided as a Source Data file.

### 2 Supplementary Tables

**Supplementary Table 1:** Results of the multilevel modeling (MLM) analysis reflecting main session effects for systems recruitment or integration (4 sessions). In all cases, random intercepts were estimated. The significance of models was estimated with chi-square tests, where models with increasing complexity were compared and the resulting value of Likelihood Ratio Test (*χ*^2^) and corresponding *p*-value (uncorrected and FDR-corrected) were reported^3^. Abbreviations: auditory (AU), cerebellum (CER), cingulo-opercular (CO), default mode (DM), dorsal attention (DA), fronto-parietal (FP), memory (MEM), salience (SAL), somatomotor (SOM), subcortical (SUB), uncertain (UNC), ventral attention (VA), and visual (VIS).

| Repeated measures MLM: main effect of session |  |  |  |
| --- | --- | --- | --- |
| Systems | Statistics |  |  |
| | $\chi^2$ | $p_{uncorr.}$ | $p_{FDR}$ |
| Recruitment |  |  |  |
| SAL | 24.3038 | 0.0000 | 0.0005 |
| DM | 24.1711 | 0.0000 | 0.0005 |
| AU | 14.7401 | 0.0021 | 0.0208 |
| MEM | 9.8965 | 0.0195 | 0.1181 |
| SUB | 8.5157 | 0.0365 | 0.1747 |
| Integration with default mode (DM) system |  |  |  |
| DM-SAL | 31.3720 | 0.0000 | 0.0001 |
| DM-AU | 20.1747 | 0.0002 | 0.0024 |
| DM-FP | 19.7550 | 0.0002 | 0.0025 |
| DM-CO | 15.8257 | 0.0012 | 0.0140 |
| Integration with task-positive systems |  |  |  |
| FP-SAL | 28.8229 | 0.0000 | 0.0001 |
| DA-SAL | 21.8571 | 0.0001 | 0.0013 |
| CO-DA | 14.0292 | 0.0029 | 0.0261 |
| MEM-SOM | 13.1715 | 0.0043 | 0.0354 |
| SAL-MEM | 11.0954 | 0.0112 | 0.0785 |
| SAL-SUB | 10.0385 | 0.0182 | 0.1181 |
| FP-CO | 9.7186 | 0.0211 | 0.1201 |
| CO-VIS | 9.499441 | 0.023337 | 0.124923 |
| SAL-CO | 8.9760 | 0.0296 | 0.1497 |
| FP-AU | 8.2521 | 0.0411 | 0.1869 |
| VA-UNC | 7.9361 | 0.0474 | 0.2032 |
| Other |  |  |  |
| AU-SOM | 11.6143 | 0.0088 | 0.0669 |

**Supplementary Table 2:** Planned contrasts for all significant main session effects, reflecting changes of systems recruitment or integration (4 sessions). Contrasts: ‘Naive’ vs. ‘Early’, ‘Naive’ vs. ‘Middle’, ‘Naive’ vs. ‘Late’. Abbreviations: auditory (AU), cerebellum (CER), cingulo-opercular (CO), default mode (DM), dorsal attention (DA), fronto-parietal (FP), memory (MEM), salience (SAL), somatomotor (SOM), subcortical (SUB), uncertain (UNC), ventral attention (VA), and visual (VIS).

| **Planned contrasts: main effect of session** |  |  |  |  |  |  |
| --- | --- | --- | --- | --- | --- | --- |
| Systems | Naive vs. Early |  | Naive vs. Middle |  | Naive vs. Late |  |
| | $\beta$ | $p$ | $\beta$ | $p$ | $\beta$ | $p$ |
| Recruitment |  |  |  |  |  |  |
| SAL | 0.0259 | 0.0279 | 0.041229 | 0.0006 | 0.057214 | 0.0000 |
| DM | 0.0358 | 0.0528 | 0.052197 | 0.0051 | 0.091959 | 0.0000 |
| AU | 0.0377 | 0.0127 | 0.0314 | 0.0369 | 0.0572 | 0.0002 |
| MEM | 0.0292 | 0.1145 | 0.016409 | 0.3740 | 0.05636 | 0.0027 |
| SUB | 0.0409 | 0.0145 | 0.015256 | 0.3569 | 0.039226 | 0.0190 |
| Integration with default mode (DM) system |  |  |  |  |  |  |
| DM-SAL | -0.0378 | 0.0008 | -0.047378 | 0.0000 | -0.0623 | 0.0000 |
| DM-AU | -0.0237 | 0.0579 | -0.0174 | 0.1633 | -0.0560 | 0.0000 |
| DM-FP | -0.0116 | 0.3310 | -0.021237 | 0.0758 | -0.051636 | 0.0000 |
| DM-CO | -0.0273 | 0.0399 | -0.0323 | 0.0154 | -0.0529 | 0.0001 |
| Integration with task-positive systems |  |  |  |  |  |  |
| FP-SAL | 0.0395 | 0.0120 | 0.062749 | 0.0001 | 0.082824 | 0.0000 |
| DA-SAL | 0.0420 | 0.0009 | 0.05039 | 0.0001 | 0.051894 | 0.0001 |
| CO-DA | 0.0271 | 0.0512 | 0.0356 | 0.0109 | 0.0511 | 0.0003 |
| MEM-SOM | -0.0408 | 0.0201 | -0.030599 | 0.0798 | -0.062644 | 0.0004 |
| SAL-MEM | 0.0109 | 0.4989 | 0.015714 | 0.3298 | 0.050995 | 0.0019 |
| SAL-SUB | 0.0364 | 0.0037 | 0.029535 | 0.0179 | 0.027495 | 0.0272 |
| FP-CO | 0.0276 | 0.0736 | 0.0349 | 0.0240 | 0.0459 | 0.0032 |
| CO-VIS | 0.024047 | 0.0535 | 0.022812 | 0.0668 | 0.037705 | 0.0027 |
| SAL-CO | 0.0257 | 0.0152 | 0.0192 | 0.0683 | 0.0285 | 0.0072 |
| FP-AU | -0.0167 | 0.1912 | -0.0025 | 0.8442 | 0.0196 | 0.1242 |
| VA-UNC | -0.0103 | 0.1671 | -0.020728 | 0.0059 | -0.008145 | 0.2726 |
| Other |  |  |  |  |  |  |
| AU-SOM | 0.0407 | 0.0056 | 0.0337 | 0.0213 | 0.0446 | 0.0025 |

**Supplementary Table 3:** Results of the multilevel modeling (MLM) analysis reflecting session × group interaction effects for systems recruitment or integration (4 sessions, 2 groups). In all cases, random intercepts were estimated. The significance of models was estimated with chi-square tests, where models with increasing complexity were compared and the resulting value of Likelihood Ratio Test (*χ*^2^) and corresponding *p*-value (uncorrected and FDR-corrected) were reported^3^. Abbreviations: auditory (AU), cerebellum (CER), cingulo-opercular (CO), default mode (DM), dorsal attention (DA), fronto-parietal (FP), memory (MEM), salience (SAL), somatomotor (SOM), subcortical (SUB), uncertain (UNC), ventral attention (VA), and visual (VIS).

| **Repeated measures MLM: session × group interaction** |  |  |  |
| --- | --- | --- | --- |
| Systems | Statistics |  |  |
| | $\chi^2$ | $p_{uncorr.}$ | $p_{FDR}$ |
| Recruitment |  |  |  |
| FP | 6.831 | 0.029 | 0.327 |
| DA | 2.697 | 0.041 | 0.339 |
| Integration with default mode (DM) system |  |  |  |
| DM-FP | 19.755 | 0.003 | 0.235 |
| Integration with subcortical (SUB) system |  |  |  |
| SUB-DM | 5.999 | 0.015 | 0.278 |
| SUB-AU | 5.843 | 0.02 | 0.309 |
| SUB-VA | 4.868 | 0.037 | 0.334 |
| SUB-DA | 1.193 | 0.026 | 0.327 |
| SUB-CO | 0.257 | 0.011 | 0.244 |
| Integration with other task-positive systems |  |  |  |
| DA-SOM | 2.514 | 0.011 | 0.244 |
| CO-MEM | 0.865 | 0.034 | 0.334 |
| CO-UNC | 2.437 | 0.005 | 0.236 |

**Supplementary Table 4:**
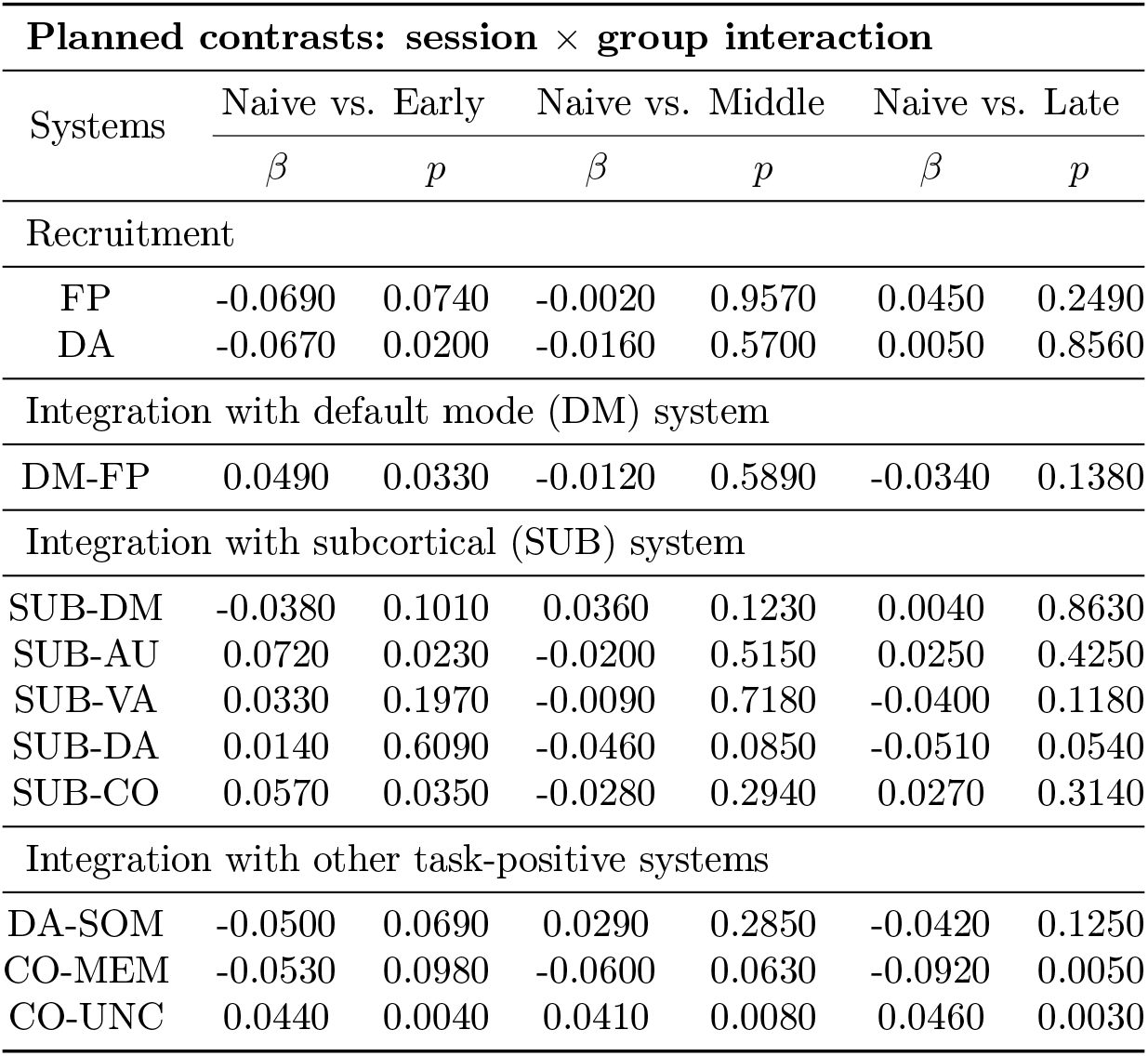
Planned contrasts for all significant session × group interaction effects, reflecting group differences in changes of systems recruitment or integration (4 sessions, 2 groups). Contrasts: ‘Naive’ vs. ‘Early’, ‘Naive’ vs. ‘Middle’, ‘Naive’ vs. ‘Late’. Abbreviations: auditory (AU), cerebellum (CER), cingulo-opercular (CO), default mode (DM), dorsal attention (DA), fronto-parietal (FP), memory (MEM), salience (SAL), somatomotor (SOM), subcortical (SUB), uncertain (UNC), ventral attention (VA), and visual (VIS).

**Supplementary Table 5:** Post-hoc tests for all significant session × group interaction effects, reflecting group differences in changes of systems recruitment or integration (4 sessions, 2 groups). Tests: ‘Naive’ vs. ‘Early’, ‘Early’ vs. ‘Middle’, ‘Middle’ vs. ‘Late’. Abbreviations: auditory (AU), cerebellum (CER), cingulo-opercular (CO), default mode (DM), dorsal attention (DA), fronto-parietal (FP), memory (MEM), salience (SAL), somatomotor (SOM), subcortical (SUB), uncertain (UNC), ventral attention (VA), and visual (VIS).

| **Post-hoc tests: session × group interaction** |  |  |  |  |
| --- | --- | --- | --- | --- |
| Systems | Early vs. Middle |  | Middle vs. Late |  |
| | $\beta$ | $p_{bonferroni}$ | $\beta$ | $p_{bonferroni}$ |
| Recruitment |  |  |  |  |
| FP | 0.0673 | 0.2499 | 0.0467 | 0.6842 |
| DA | 0.0506 | 0.2245 | 0.0212 | 1.0000 |
| Integration with default mode (DM) system |  |  |  |  |
| DM-FP | -0.0613 | 0.0237 | -0.0216 | 1.0000 |
| Integration with subcortical (SUB) system |  |  |  |  |
| SUB-DM | 0.0739 | 0.0052 | -0.0318 | 0.5121 |
| SUB-AU | -0.0919 | 0.0114 | 0.0453 | 0.4462 |
| SUB-VA | -0.0418 | 0.2997 | -0.0305 | 0.6836 |
| SUB-DA | -0.0593 | 0.0786 | -0.0055 | 1.0000 |
| SUB-CO | -0.0854 | 0.0055 | 0.0554 | 0.1229 |
| Integration with other task-positive systems |  |  |  |  |
| DA-SOM | 0.0790 | 0.0130 | -0.0711 | 0.0300 |
| CO-MEM | -0.0068 | 1.0000 | -0.0321 | 0.9536 |
| CO-UNC | -0.0033 | 1.0000 | 0.0054 | 1.0000 |

**Supplementary Table 6:** Correlations between the change in network dynamics and the change in behavior. Pearson correlation coefficient (*r*) between the across-session changes (Naive vs. Late) in recruitment (or integration) and the across-session changes in *d*′ (Δ*d*′) and pRT (Δ pRT) observed for both the experimental and control groups. Abbreviations: auditory (AU), cerebellum (CER), cingulo-opercular (CO), default mode (DM), dorsal attention (DA), fronto-parietal (FP), memory (MEM), salience (SAL), somatomotor (SOM), subcortical (SUB), uncertain (UNC), ventral attention (VA), and visual (VIS).

| **Correlation with behavior: Naive vs. Late (both groups)** |  |  |  |
| --- | --- | --- | --- |
| Systems | $r$ | $p_{uncorr.}$ | $p_{FDR}$ |
| $\Delta d'$ and $\Delta$ recruitment | | | |
| SAL | 0.3377095 | 0.028723065 | 1 |
| DM | 0.3354091 | 0.029898227 | 1 |
| $\Delta d'$ and $\Delta$ integration | | | |
| FP-SAL | 0.3530235 | 0.021836677 | 1 |
| MEM-VIS | 0.3215687 | 0.037836231 | 1 |
| SOM-VA | 0.3198655 | 0.038922327 | 1 |
| DM-FP | -0.3099325 | 0.04577489 | 1 |
| DM-SAL | -0.4142908 | 0.006378749 | 0.5740874 |
| AU-MEM | -0.4412674 | 0.003442311 | 0.3132503 |
| $\Delta$ pRT and $\Delta$ recruitment | | | |
| DM | -0.3478555 | 0.02398683 | 1 |
| VIS | -0.3378605 | 0.02864729 | 1 |
| $\Delta$ pRT and $\Delta$ integration | | | |
| AU-MEM | 0.3178699 | 0.04022718 | 1 |
| CER-DM | 0.3405356 | 0.02733201 | 1 |
| CO-UNC | -0.3630498 | 0.01812347 | 1 |
| CO-VIS | 0.3338558 | 0.03071395 | 1 |
| DM-FP | 0.3560102 | 0.02066934 | 1 |

**Supplementary Table 7:** Relationship between the change in network dynamics and the change in behavior. Pearson correlation coefficient (*r*) between the changes in recruitment (or integration) and the changes in *d*′ (Δ*d*′) and pRT (Δ pRT) during early phase of training (Naive vs. Early) of the experimental group. Abbreviations: auditory (AU), cerebellum (CER), cingulo-opercular (CO), default mode (DM), dorsal attention (DA), fronto-parietal (FP), memory (MEM), salience (SAL), somatomotor (SOM), subcortical (SUB), uncertain (UNC), ventral attention (VA), and visual (VIS).

| **Correlation with behavior: Naive vs. Early (experimental)** |  |  |  |
| --- | --- | --- | --- |
| Systems | $r$ | $p_{uncorr.}$ | $p_{FDR}$ |
| $\Delta d'$ and $\Delta$ integration | | | |
| AU-VA | 0.4903851 | 0.0240127927 | 1 |
| CER-DA | 0.4585068 | 0.0365743723 | 1 |
| CO-DA | 0.4957494 | 0.0222872683 | 1 |
| CO-SUB | -0.515905 | 0.0166670006 | 1 |
| DA-SOM | 0.710518 | 0.0003068366 | 0.02792213 |
| DA-SUB | 0.4700412 | 0.0315444066 | 1 |
| DM-SAL | 0.4558561 | 0.037814337 | 1 |
| FP-SOM | 0.449137 | 0.0411045974 | 1 |
| $\Delta$ pRT and $\Delta$ integration | | | |
| AU-VA | -0.4498745 | 0.04073297 | 1 |
| CER-DA | -0.4463136 | 0.042551842 | 1 |
| CO-DA | -0.4645735 | 0.033856106 | 1 |
| CO-SUB | 0.5106186 | 0.018016262 | 1 |
| DA-SOM | -0.5641078 | 0.007729913 | 0.7034221 |
| UNC-VA | 0.5189307 | 0.015932156 | 1 |

**Supplementary Table 8:** Results of the multilevel modeling (MLM) analysis reflecting session × group interaction effects for systems activity estimated with a standard GLM (2-back vs. 1-back contrast, two-sided). In all cases, random intercepts were estimated. The significance of models was estimated with chi-square tests, where models with increasing complexity were compared and the resulting value of Likelihood Ratio Test (*χ*^2^) and corresponding p-value (uncorrected and FDR-corrected) were reported^3^. Abbreviations: auditory (AU), cerebellum (CER), cingulo-opercular (CO), default mode (DM), dorsal attention (DA), fronto-parietal (FP), memory (MEM), salience (SAL), somatomotor (SOM), subcortical (SUB), uncertain (UNC), ventral attention (VA), and visual (VIS).

| Repeated measures MLM: session × group interaction (GLM) | | | |
| --- | --- | --- | --- |
| System | Statistics |  |  |
| | $\chi^2$ | $p_{uncorr.}$ | $p_{FDR}$ |
| SAL | 19.289622 | 0.000238 | 0.003096 |
| VIS | 12.21532 | 0.006681 | 0.043425 |
| DM | 10.220241 | 0.016784 | 0.060016 |
| UNC | 9.538138 | 0.022929 | 0.060016 |
| FP | 9.523477 | 0.023083 | 0.060016 |
| MEM | 8.635576 | 0.03455 | 0.074858 |
| VA | 7.979876 | 0.046429 | 0.086226 |
| DA | 5.976738 | 0.112747 | 0.183215 |
| SOM | 4.77535 | 0.189006 | 0.273008 |
| CO | 4.233999 | 0.23728 | 0.298229 |
| SUB | 4.085796 | 0.252347 | 0.298229 |
| CER | 3.326098 | 0.344027 | 0.372696 |
| AU | 2.794743 | 0.424366 | 0.424366 |

**Supplementary Table 9:** Planned contrasts for all significant session × group interaction effects, reflecting group differences in changes of systems activity estimated with a standard GLM (2-back vs. 1-back contrast, two-sided). Contrasts: ‘Naive’ vs. ‘Early’, ‘Naive’ vs. ‘Middle’, ‘Naive’ vs. ‘Late’. Abbreviations: auditory (AU), cerebellum (CER), cingulo-opercular (CO), default mode (DM), dorsal attention (DA), fronto-parietal (FP), memory (MEM), salience (SAL), somatomotor (SOM), subcortical (SUB), uncertain (UNC), ventral attention (VA), and visual (VIS).

| **Planned contrasts: session × group interaction effect (GLM)** |  |  |  |  |  |  |
| --- | --- | --- | --- | --- | --- | --- |
| System | Naive vs. Early |  | Naive vs. Middle |  | Naive vs. Late |  |
| | $\beta$ | $p$ | $\beta$ | $p$ | $\beta$ | $p$ |
| SAL | -1.09808 | 0.000812 | -1.317647 | 0.000069 | -0.956718 | 0.003351 |
| VIS | 0.121433 | 0.684265 | -0.80268 | 0.00806 | -0.002907 | 0.992229 |
| DM | -0.750571 | 0.01187 | -0.759561 | 0.010918 | -0.779999 | 0.009004 |
| UNC | -0.255562 | 0.27055 | -0.689108 | 0.003441 | -0.206578 | 0.372717 |
| FP | -0.679019 | 0.095265 | -1.235466 | 0.002735 | -0.695647 | 0.087521 |
| MEM | -0.574095 | 0.111209 | -1.02874 | 0.004776 | -0.401063 | 0.264535 |
| VA | -0.404449 | 0.195934 | -0.76255 | 0.01565 | -0.732185 | 0.020184 |

### 3 Supplementary Methods

#### 3.1 Penalized reaction time calculation

To measure behavioral performance in the dual n-back scanning sessions, we incorporated *penalized reaction time* (pRT), which is a measure previously introduced by^4^. This measure combines both measures of accuracy and response time. For every subject, session, task condition, and stimulus modality (auditory, spatial), pRT was defined as:

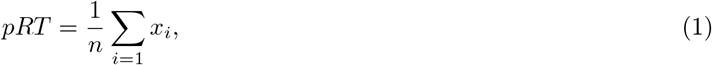

where *n* is the sum of all subject responses and incorrect response omissions, and *x_i_* was obtained from the following formula:

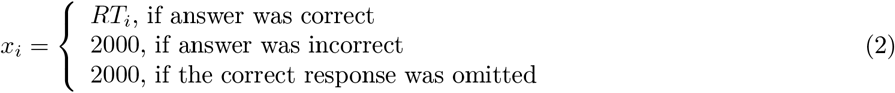

where *RT_i_* is reaction time of the response during the *i*-th trial and the scalar value of 2000 is a penalty for an incorrect answer or for the lack of an answer, which is the maximum possible time to respond during each n-back trial measured in milliseconds. For each participant, we calculated average pRT for both modalities to represent a cumulative measure of performance during the dual n-back task.

#### 3.2 Behavioral variability analysis

To assess measures of behavioral variability, we calculated (1) block-wise variants of the two behavioral performance measures, d’ and penalized reaction time (pRT), and (2) the standard deviation of these measures over task blocks. For consistency with the measures used in the main text, for both block-wise measures we considered the average value over both stimulus modalities (visual and auditory). This procedure resulted in two measures of block-to-block behavioral variability for each participant and session: the standard deviation of d’ (*σ*_*d*′_) and the standard deviation of pRT (*σ_pRT_*). We then used a multilevel analysis to investigate group × session interactions. Note that these measures of behavioral variability can potentially capture two distinct effects: (1) more or less consistent performance during the 1-back or 2-back blocks, and (2) greater or lesser decreases in behavioral performance from the 1-back to the 2-back condition. Both effects of more consistent performance during a single task condition and a lesser decrease in performance from the 1-back to the 2-back condition would result in an overall decrease in the behavioral variability measures of *σ*_*d*′_ and *σ_pRT_*.

#### 3.3 Anatomical data processing in fMRIPrep

Each T1w (T1-weighted) volume was corrected for INU (intensity non-uniformity) using N4BiasFieldCorrection v2.1.0^5^ and skull-stripped using antsBrainExtraction.sh v2.1.0 (employing the OASIS template). Brain surfaces were reconstructed using recon-all from FreeSurfer v6.0.1^6^, and the brain mask estimated previously was refined with a custom variation of the method to reconcile ANTs-derived and FreeSurfer-derived segmentations of the cortical gray-matter of Mindboggle^7^. Spatial normalization to the ICBM 152 Nonlinear Asymmetrical template version 2009c^8^ was performed through nonlinear registration with the antsRegistration tool of ANTs v2.1.0^9^, using brain-extracted versions of both the T1w volume and template. Brain tissue segmentation of cerebrospinal fluid (CSF),white matter (WM), and gray matter was performed on the brain-extracted T1w using FAST^10^ (FSL v5.0.9).

#### 3.4 Multilevel community detection for signed networks

We ran multilayer community detection on networks with both positive and negative edges^11,12^, to investigate whether the antagonism between large-scale systems (reflected by anticorrlated time-series) could influence the recruitment and integration values. First, we defined *N* × *N* matrix 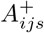 by zeroing negative elements of *A_ijs_* and *N* × *N* matrix 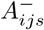 by zeroing positive elements of *A_ijs_*. We used this decomposition to represent both *A_ijs_* and the corresponding null model *p_ijs_* as a linear combination of networks with positive and networks with negative edges:

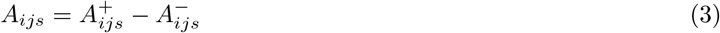

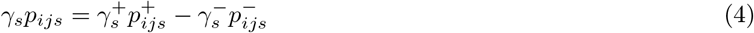

Then, we maximized following modularity quality function:

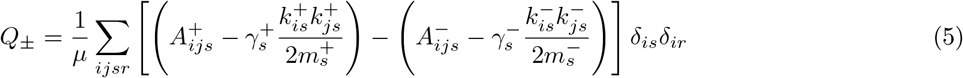

With this approach we consider the negative network edges as separate networks when calculating within-layer modularity.

#### 3.5 Standard GLM analysis

To enable reference to the prior literature on the effects of working memory training on activation patterns, we additionally performed a standard General Linear Model (GLM) analysis. In the first level of the GLM analysis, we compared 2-back vs. 1-back activation patterns (two-sided) for all subjects to identify brain areas activated and deactivated in a more difficult 2-back condition. Then, we ran a second-level GLM analysis to investigate consistent patterns of task activation in all sessions and both groups. To make GLM analysis comparable with our functional connectivity analysis, we calculated the mean z-score for the first-level /*beta* maps for each ROI from the Power et al.^1^ parcellation (Supplementary Figure 10). Then, for all large-scale systems we calculated the mean z-score that reflected the effect size for each network, and sorted them from the lowest to the highest. Next, we used multilevel modelling to test for session × group interactions for each system (see Supplementary Figure 11 and Supplementary Table 8–9).

## Notes

#### Summary of Updates

Manuscript text updated; Supplemental files updated.

https://osf.io/wf85u/

